# Abiotic conditions shape spatial and temporal morphological variation in North American birds

**DOI:** 10.1101/2022.02.17.480905

**Authors:** Casey Youngflesh, James F. Saracco, Rodney B. Siegel, Morgan W. Tingley

## Abstract

Abiotic environmental conditions play a key role in driving the size and shape of organisms. Quantifying environment-morphology relationships is important not only for understanding the fundamental processes driving phenotypic diversity within and among species (*1*), but also for predicting how species will respond to ongoing global change (*2*). Despite a clear set of expectations motivated by ecological theory (*3*), broad evidence in support of generalizable effects of abiotic conditions, such as temperature (*4*), on spatial and temporal intraspecific morphological variation has been limited. Using standardized data from over 250,000 captures of 105 landbird species, we assessed intraspecific shifts in bird morphology since 1989 while simultaneously measuring spatial morphological gradients across the North American continent. Across bird species, we found strong spatial and temporal trends in body size, with warmer temperatures associated with smaller body sizes both at more equatorial latitudes and in more recent years. The magnitude of these thermal effects varied both across and within species, with results suggesting it is the warmest, rather than the coldest, temperatures driving both spatial and temporal trends. Across elevation, we found that body size declines as relative wing length increases, likely due to the benefits that longer wings confer for flight in thin air environments. Our results provide support for both existing and new large-scale ecomorphological gradients and highlight how the response of functional tradeoffs to abiotic variation drives morphological change.

**Significance Statement:** Characterizing how the size and shape of organisms varies over space and time is key to understanding the processes that create ecological communities and for predicting how species will respond to climate change. Across more than 100 species of North American birds, we show that within species the size and shape of individuals varies substantially across space and time. Warmer temperatures are associated with smaller body sizes, likely due to the importance of body size for thermoregulation. As the climate continues to warm, these species will likely continue to shrink. We also provide the first large-scale evidence of an increase in wing length with elevation, a pattern that could be attributed to thinner air in high elevation environments.

## INTRODUCTION

Morphology is both a cause (*5*) and a consequence (*6*) of how organisms interact with their environment. Assessing patterns in morphological variation both across and within (*7*) species provides a means to better understand these interactions, and consequently, predict ecological responses to environmental change. Ecological theory suggests that both the sizes and shapes of organisms should vary across latitude [e.g., Bergmann’s (*3*) and Allen’s (*8*) Rules] and also possibly elevation, particularly for flying organisms [due to lower temperatures and lower air density at high elevations (*9*)]. These ecogeographic expectations are commonly used to motivate hypotheses for how species will respond to climate change (*10*), such as the suggestion that declining body size may be a generalized response of endotherms to warming temperatures (*2*). However, broad evidence in support of generalizable effects of abiotic conditions on intraspecific spatial and temporal morphological variation has been limited by a lack of taxonomic and spatial replication (*2, 11–13*), yielding conflicting results. For birds in particular, which have precipitously declined in North America over a period coincident with modern anthropogenic warming (*14*), much remains unknown regarding how abiotic factors may shape morphological traits over space and time.

We evaluated spatiotemporal morphological variation in 105 North American bird species over 30 years (1989–2018), across more than 43 degrees of latitude and nearly 3,000 m of elevation, using data from more than 250,000 live birds, primarily passerines or near-passerines, captured during the breeding season using standardized methods (*15*) (Fig. 1A, S1, Table S1). These measurements comprise a dataset on bird morphology that is unparalleled in size, taxonomic diversity, and spatiotemporal scope. Combining field measures of body mass and wing length (i.e., length of the unflattened, closed wing) with allometric scaling theory (*16*), we derived two morphological indices, a Size Index (*SI*) and a Wing Index (*WI*)(Fig. 1B, 1C). *SI* and *WI* reflect overall bird body size and “wingyness” (wing length relative to body mass), respectively (Fig. S2), and were used to account for the fact that mass and wing length are intrinsically linked (i.e., that changes in mass may be due to changes in wing length and vice versa). Using a hierarchical Bayesian approach to estimate species-specific responses, we modeled these indices as a function of year, latitude, and elevation, and estimated the impact of spatial and temporal variation in temperature on bird body size.

**Fig. 1.**
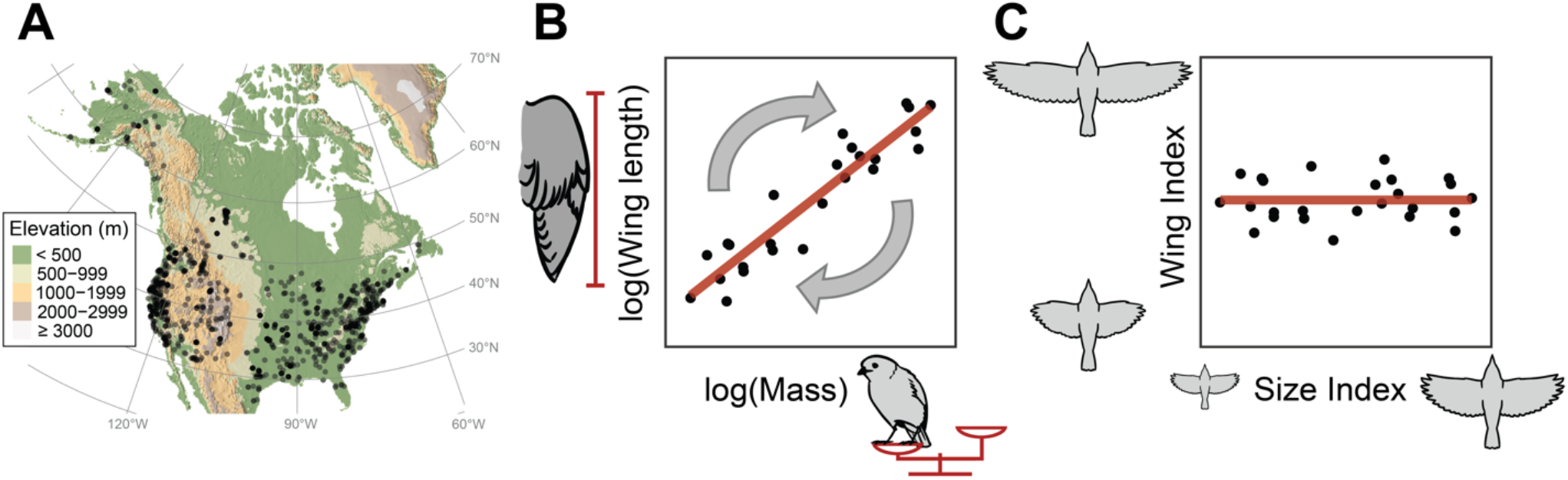
The Monitoring Avian Productivity and Survivorship (MAPS) dataset provides an unparalleled resource for studying avian morphological variation over time and space. (**A**) Data on individual birds come from 1124 MAPS banding stations (black points) spanning the latitudinal and elevational extent of North America. (**B**) Measurements were taken for both wing length (chord of the unflattened wing) and mass for each captured bird. Based on allometric scaling principals and empirical measurements across species, wing length is expected to be proportional to mass to the 1/3 power (the scaling exponent in the power law equation); logging both variables linearizes this relationship. Points represent individuals from a single hypothetical species. (**C**) The scaling exponent was used to create a rotation matrix which was applied to logged wing length and logged mass for each species, to derive two independent morphological indices: a Size Index (*SI*) and a Wing Index (*WI*), denoting the overall size of each individual bird and the degree to which wing length deviates from its expected value given the body mass of the individual, respectively. For additional details on this mathematical transformation, see Fig. S2.

## RESULTS

Across the wide spatial and taxonomic breadth of sampling, avian body size decreased over time (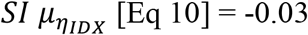 *SI* per 10 years, 89% CI: [−0.04, −0.01], *p*(< 0) = 1; Fig. 2A, S3A, Table S2). Absolute body mass showed range-wide declines of up to 2.78% between 1989 and 2018 (e.g., tree swallow *Tachycineta bicolor*, 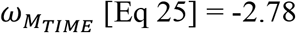, 89% CI: [−4.98, −0.63], *p*(< 0) = 0.98), with a mean decline in mass of 0.56% across all 105 species 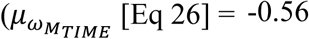, 89% CI: [−0.78, −0.34], *p*(< 0) = 1; Fig. S7A, Table S3). This temporal trend toward smaller bodies, replicated across species and over most of a continent, is likely the result of warming summer temperatures. Specifically, smaller body sizes were associated with elevated June maximum temperatures in the year of capture (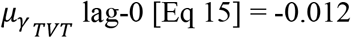 *SI* per 1° C, 89% CI: [−0.014, −0.009], *p*(< 0) = 1), as well as one year prior to capture (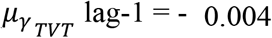, 89% CI: [−0.007, −0.001], *p*(< 0) = 0.99). Temperatures two years prior to capture were not strongly related to body size (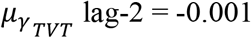, 89% CI: [−0.003, 0.001], *p*(< 0) = 0.73; Fig. 2B, S5, Table S4). Temperatures one and two years prior to capture correspond to environmental conditions likely experienced during ontogenesis, although post-natal dispersal limits the strength of this inference from banding data. However, these findings align with expectations, given that smaller-bodied individuals – having larger surface-area-to-volume ratios– tend to have lower cooling costs compared to larger-bodied individuals, and provide strong support for the hypothesis that shrinking body size is a generalized response to climate change (*2, 10*).

**Fig. 2:**
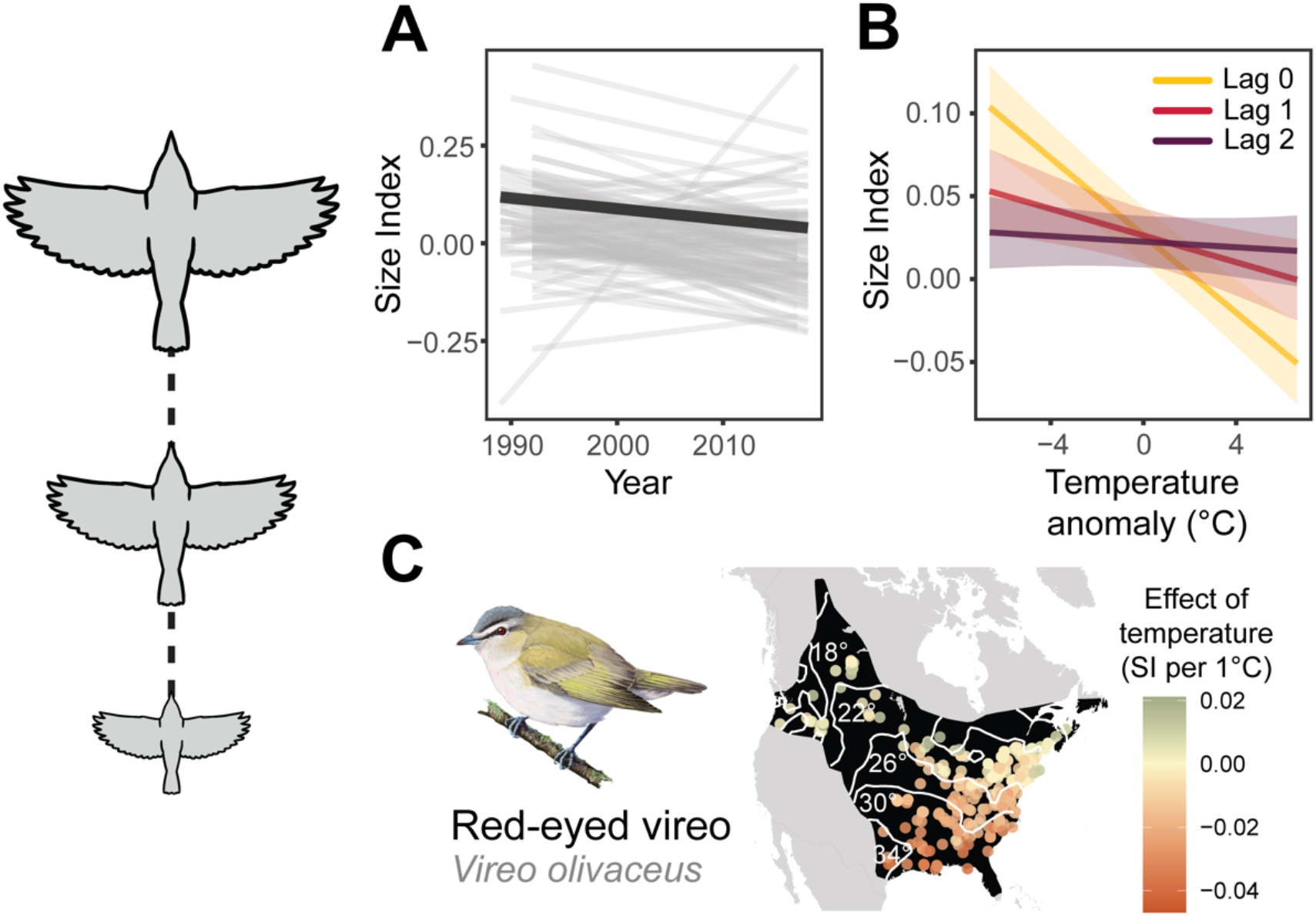
Change in Size Index (*SI*) over time and in response to temporal fluctuations in temperature. (**A**) Change in *SI* over time for 105 species, controlling for changes over latitude and elevation. Each thin gray line represents the trend for one species, while the thick black line represents the mean trend across all species. (**B**) Change in *SI* across species in response to inter-annual fluctuations in June maximum temperature in the year of capture (Lag 0) as well as one (Lag 1) and two years (Lag 2) prior to capture. Ribbons represent 89% CIs. (**C**) Effect of 1°C change in temperature on *SI* at capture locations for a representative species, the red-eyed vireo *Vireo olivaceus*, showing stronger effects of temperature on *SI* in warmer areas. Darker, orange hues represent a stronger negative effect of temperature on *SI*. The black polygon represents the range of the species, while white lines (and associated white text) represent isoclines for June maximum temperature in a single year, 2018.

Temperature-mediated size-dependent mortality [which may result in directional selection, conditional on the heritability of body size; e.g., (*17*)] and/or developmental plasticity during early life stages (*18*) may be the most likely proximate drivers of our finding of an association between warmer summers and smaller bodies. While widespread evidence for adaptive evolutionary responses to climate change is somewhat limited (*19*), the rate of morphological change reported here is within the range that might be expected via evolutionary change (Fig. S10). The fact that conditions in the year of capture have the strongest effect on body size may indicate that temperature impacts morphology most strongly via size-dependent mortality of adult birds. The lack of a strong relationship with temperatures two years prior to capture could suggest that a large portion of measured individuals were in their second year of life and never experienced the conditions 24 months prior. Greater effects of temperature on body size in the warmer portions of species’ ranges (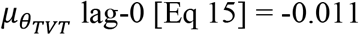 unit change in effect of temperature per 10° C change in mean site temperature, 89% CI: [−0.020, −0.002], *p*(< 0) = 0.97; Fig. 2C) suggests that it is the hottest experienced temperatures – rather than the coldest – driving this body size-temperature association (*20*). This effect was less pronounced for temperatures in the year prior and two years prior to capture (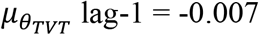, [89% CI: −0.017, 0.002], *p*(< 0) = 0.90; 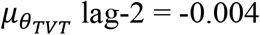, 89% CI: [−0.014, 0.005], *p*(< 0) = 0.75; Fig. 2C). While poleward range shifts of species could also result in directional change in morphology at a given location, declines in body size in even the warmest portions of species’ ranges (where individuals are generally smallest) suggests that dispersal is not the primary mechanism driving these observed changes.

In contrast to shrinking body size in North American birds, we found that the wingyness (wing length relative to body mass) of birds has increased over time (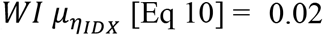 *WI* per 10 years, 89% CI: [0.00, 0.03], *p*(> 0) = 0.95; Fig. S4A, Table S2). While this pattern could be due to changing migratory patterns in response to ongoing range shifts (*21*), constraints on the rate at which wing length can change over time compared to body size (*22, 23*) might also play a role. Specifically, in contrast to previous findings that relied on bird specimens derived from a single migratory bottleneck (*13*), we observed no change in absolute wing length over time – temporal changes in wingyness were the result of declining mass (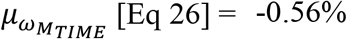 change over study period, 89% CI: [−0.78, −0.34], *p*(< 0) = 1) with relatively stable wing length (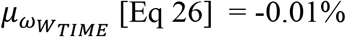 change over study period, 89% CI: [−0.13, 0.12], *p*(< 0) = 0.54; Fig. S7A, S8A, Table S3). That is, while birds have overall gotten smaller, their wings have stayed relatively the same size.

Why is it so critical to control for geography when assessing temporal trends in phenotypes? Bird morphology shows strong and generalizable trends in morphology over both latitude and elevation. As illustrated by our dataset across 105 bird species and most of a continent, body size strongly increases with latitude (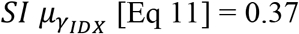 *SI* per 10° latitude, 89% CI: [0.29, 0.45], *p*(> 0) = 1; Fig. 3A, S3B, Table S2), supporting the intraspecific interpretation of Bergmann’s Rule (*3*), despite decades of debate on its relevance (*4*). Across species’ latitudinal ranges, body mass increases 5.72% (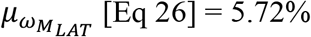, 89% CI: [5.39, 6.04], *p* > 0) = 1; Fig. S7B, Table S3). Larger body sizes are associated with regions with cooler average temperatures (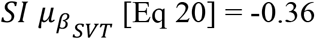 *SI* per 10° C change in mean site temperature, 89% CI: [−0.44, −0.27], *p*(< 0) = 1; Fig. 3B, S6) that are generally found at higher latitudes, supporting the notion that thermal factors play a substantial role in governing body size not only over time, but also over space (*20*). Additionally, we found that this relationship between temperature and spatial variation in body size is stronger for species that experience warmer conditions (*θ*_*SR*_ [Eq 22] = −0.30 unit change in effect of temperature per 10° C change in mean range-wide temperature, 89% CI: [−0.49, −0.10], *p*(< 0) = 0.99; Fig. 3B), illustrating – as with findings of temporal associations between body size and temperature – that the warmest, rather than the coldest, temperatures likely drive intraspecific adherence to Bergmann’s Rule.

**Fig. 3:**
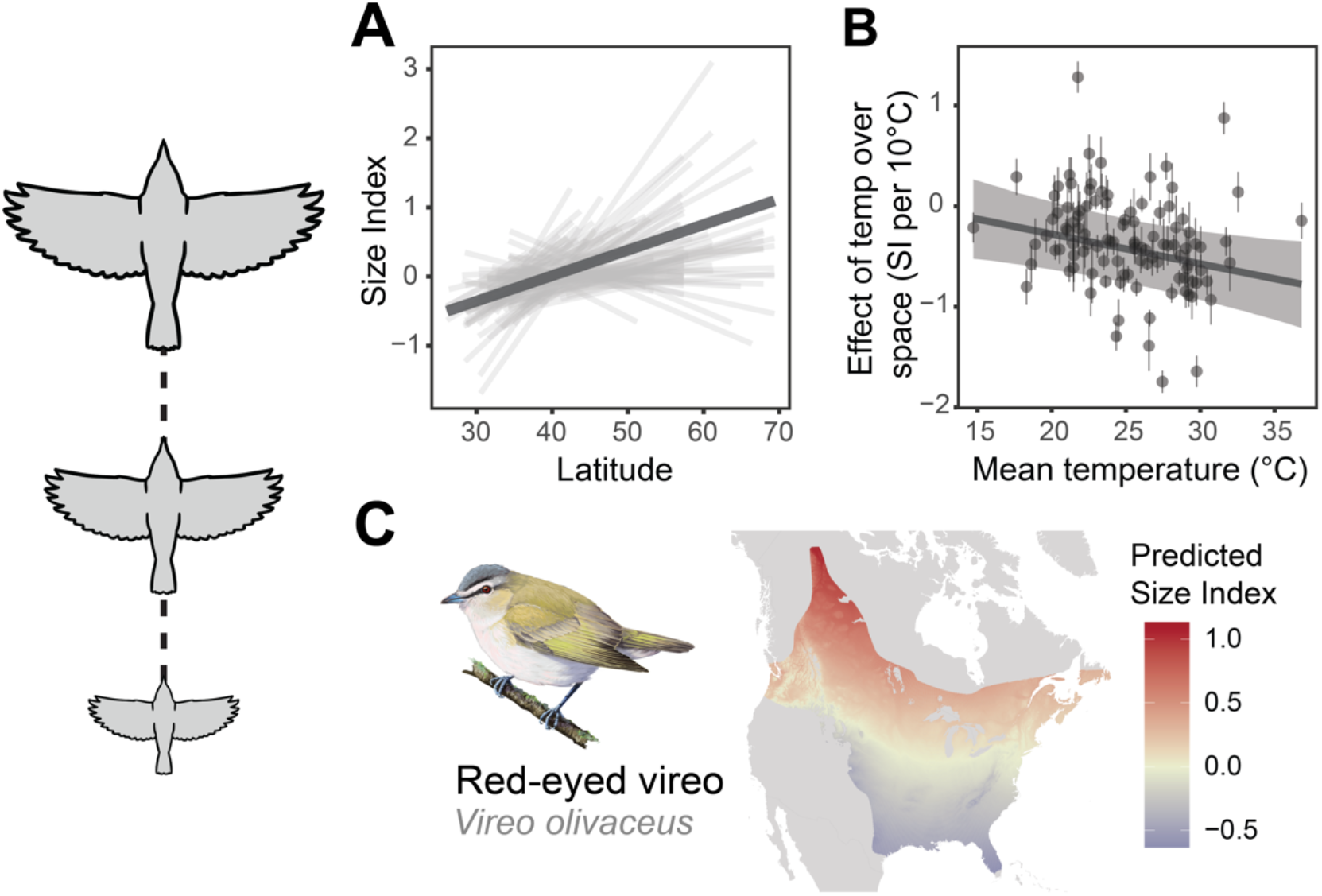
Change in *SI* over latitude and in response to spatial variation in temperature. (**A**) Change in *SI* over latitude for 105 species, controlling for changes over time and elevation. Each thin gray line represents the trend for one species, while the thick black line represents the mean trend across all species. (**B**) The effect of spatial variation in temperature on *SI* within each species as a function of the mean (range-wide) temperature experienced by that species. Each point represents a single species. Gray vertical bars represent one posterior standard deviation of the effect of spatial variation in temperature on *SI*, the thick black line represents the linear model fit, and the gray ribbon represents the 89% CI. (**C**) Predicted body size (*SI*) over the range of a representative species, the red-eyed vireo *Vireo olivaceus*, based on the estimated effect of latitude and elevation. Yellow hues represent average, red hues represent larger than average, and blue hues represent smaller than average predicted *SI*.

Factors other than temperature may also be important in driving morphological variation. For example, some evidence exists for an increase in wingyness with latitude (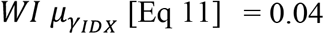 *WI* per 10° latitude, 89% CI: [0.00, 0.07], *p*(> 0) = 0.92; Fig. S4B, Table S2). While thermal factors might suggest that appendages should be smaller towards the poles to limit heat loss – known as Allen’s rule (*8*) – bird wings are not thermoregulatory organs. The length of the closed bird wing is primarily a function of flight feather length. Relatively longer wings at higher latitudes may reflect the longer distances that breeding birds from more northerly populations tend to travel to complete their migration. Longer and more pointed wings are thought to increase the efficiency of long flights and are generally found in populations that migrate longer distances (*24*). For some species, populations breeding at higher latitudes migrate farther than southern populations, yielding ‘leapfrog’ migration patterns; for other species, equatorward populations of an otherwise migratory species remain non-migratory (*25*). Indeed, species known to exhibit leapfrog migrations [e.g., Wilson’s warbler *Cardellina pusilla*: (*26*), fox sparrow *Passerella iliaca*: (*27*)], as well as migratory species with resident populations in the southern portions of their ranges [e.g., Eastern towhee *Pipilo erythrophthalmus*: (*28*), white-eyed vireo *Vireo griseus*: (*28*)], here show pronounced increases in wingyness with latitude (Fig. S4B, Table S2). Smaller or even negative effects of latitude for other species might be indicative of alternative migration strategies – in which northerly populations do not migrate longer distances than southerly populations (*25*) – as well as the importance of other factors, such as variation in habitat structure (*29*) and/or predation (*30*), that might also drive variation in wing length.

Less well understood is how morphology varies over elevation. Given decreasing temperatures at high elevations, body size might be expected to increase (i.e., Bergmann’s Rule applied to elevation). However, we find that body size generally decreases with elevation (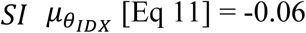 *SI* per 1000m, 89% CI: [−0.12, 0.00], *p*(< 0) = 0.96; Fig. S3C, Table S2), indicating that, contrary to the general associations found between body size and temperature over space, pressures unrelated to thermoregulation dominate over this gradient [potentially reflecting lower resource availability at higher elevations (*31*)]. Species with wide elevational gradients may therefore rely on a variety of behavioral adaptations, including altitudinal migration (*32*) and even nightly torpor (*33*), to cope with lower temperatures at higher elevations.

In contrast to body size, wingyness strongly increases with elevation (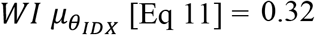 *WI* per 1000m, 89% CI: [0.28, 0.37], *p*(> 0) = 1; Fig. 4, S4C, Table S2). Elevational trends in both indices are due to countervailing changes in absolute morphology: body mass decreases (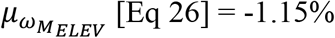 change over species’ elevational range, 89% CI: [−1.42, −0.89], *p*(< 0) = 1; Fig. S7C, Table S3) while wing length increases with elevation (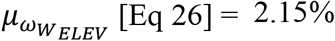 change, 89% CI: [2.00, 2.30], *p*(> 0) = 1; Fig. S8C, Table S3). These elevational ecogeographic relationships for birds are likely due to the key role that air pressure plays in flight performance. Air density, a key determinant in the amount of lift that a wing produces, is lower at higher elevations, necessitating some compensatory measures to maintain flight [i.e., more relative power output via larger wings and/or lower mass, larger wing stroke amplitude, or increased wingbeat frequency (*9, 34*)].

**Fig. 4:**
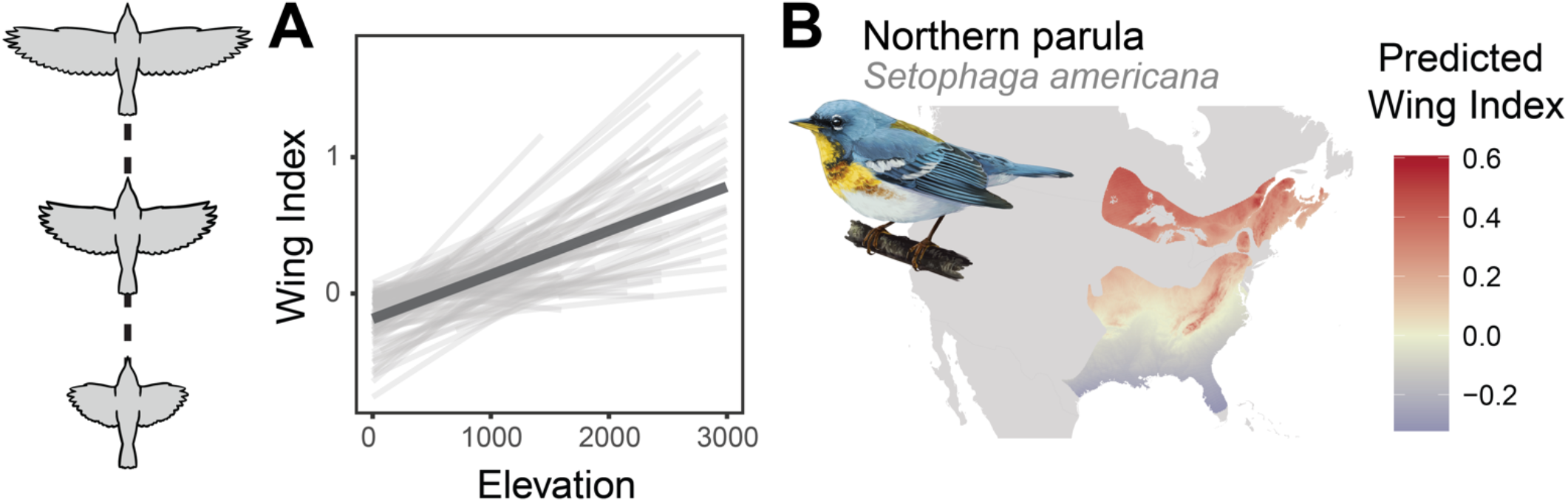
Change in *WI* over elevation and across latitude. **(A**) Change in *WI* over elevation for 105 species, controlling for changes over time and latitude. Each thin gray line represents the trend for one species, while the thick black line represents the mean trend across all species. (**B**) Predicted wingyness (*WI*) over the range of a representative species, the northern parula *Setophaga americana*, based on the estimated effect of latitude and elevation. Yellow hues represent average, red hues represent larger than average, and blue hues represent smaller than average predicted *WI*.

While large-scale increases in wing size with elevation have been documented previously, this pattern was (incorrectly) taken to be indicative of an increase in the size of individuals (*35*). Our results illustrate a clear increase in wing length with elevation independent of any changes in body size (Fig. 4), providing large-scale, cross-taxonomic evidence for a new ecomorphological gradient. This intraspecific pattern of increased wing length with elevation harmonizes observations in some insects (*36*), among specific groups of bird species, including hummingbirds [family Trochilidae (*37*)] and white-eyes [*Zosterops* spp. (*34*)], and from a limited number of single-species studies [e.g., song sparrow *Melospiza melodia* (*38*), Eurasian tree sparrow *Passer montanus* (*39*)].

## DISCUSSION

While intraspecific morphological differences are often disregarded in macroecological and functional studies [e.g., (*40*)], this important element of biodiversity has important implications for understanding how organisms are shaped by their environments, how they are likely to respond to future global change, and for the conservation of natural systems (*41*). For example, the degree to which species can respond to the thermoregulatory pressures caused by warming temperatures may impact their ability to persist in their current ranges (*42*). More frequent extreme weather events that may result in large-scale thermoregulatory-related mortality events (*43*) and chronic sub-lethal effects of increased temperature may have pronounced effects on populations (*44*). While body size in North American birds has responded to warming temperatures over time, larger responses to temperature variation over space compared to temperature variation over time (change in *SI* per 1° C was more than three times as large over space compared to temperatures over time at Lag 0) suggests that the rate of morphological change over time may be evolutionarily and/or plastically constrained – species may not be responding rapidly enough over time to keep pace with ongoing climatic change (*45*). The potential for mismatch between species and their environments is especially concerning for some bird species – such as those found in desert environments – that may lack suitable microrefugia to buffer them from especially warm temperatures (*46*).

Morphological responses to thermoregulatory pressures, as well as the importance of flight efficiency, illustrate how interacting functional tradeoffs contribute to observed morphological variation. Other factors not directly considered in this study, including conditions experienced on overwintering grounds, likely act in concert with these processes. Characterizing the interplay between these various factors, operating over space and time, is key to understanding how morphology is likely to change into the future, in response to continued abiotic environmental change. Although the ecological consequences of morphological change and how morphology interacts with other climate change responses – including shifts in species’ ranges (*47*) and the timing of seasonal events (*48*) – are currently unknown (*2*), the importance of body size for life history traits (*49*), physiology (*50*), and both cross- (*51*) and intra-trophic (*52*) interactions, suggests the implications of these changes could be far reaching. Given projected changes in climatic conditions, continued morphological change and its associated consequences can be expected into the foreseeable future.

## MATERIALS AND METHODS

### Morphological data

Bird morphology data were collected as part of the Monitoring Avian Productivity and Survivorship (MAPS) program, a collaborative long-term bird-banding project operating across North America (*15*). Data were obtained from 1124 banding stations (Fig. 1), each consisting of 6–20 mist nets, over the period 1989–2018 (though most stations operated during only a subset of this period). Banding stations were operated 6–12 times per year, from May 1 to August 28 (*15*), encompassing the breeding season for most birds in North America. Only records obtained within species’ breeding ranges were used (as determined annually by banding station operators). For each captured bird, wing length (distance between the carpal joint and the wing tip, commonly referred to as unflattened wing chord) was measured to the nearest millimeter following (*53*) and body mass was recorded to the nearest 0.5 grams (*15*). Birds were aged following criteria summarized by (*53*).

We restricted our analyses to male birds classified as After Hatch Year (captured at least one breeding season after the hatch year of the bird) to avoid any confounding morphological variation among age classes and between sexes and changes in female bird mass throughout the season that may be due to egg production and laying. All records with body mass or wing length measurements that were more than five median absolute deviations [MAD (*54*)] away from the median were excluded, as these likely represented measurement or data entry errors. If an individual was captured more than once in a season, only measurements taken during the initial capture were considered. Only species for which data were available for at least 375 captures (post data filtering) were analyzed. In total, morphological data from 253,488 captures of 105 species, representing two orders and 18 taxonomic families were used from banding stations spanning more than 43 degrees of latitude (26.1°N – 69.4°N) and 2996 meters of elevation (Table S1).

### Elevation Data

Elevation data for each banding station were obtained from the 30-arcsecond resolution (approximately 1-km at the equator) Global Multi-resolution Terrain Elevation Data 2010 (*55*) data product.

### Temperature Data

Daily maximum temperature data for each banding station were obtained over the study period from the 1-km gridded Daymet surface weather data product (*56*). For each year at each site, we calculated the average maximum temperature from ordinal day (day of year) 152 to ordinal day 181 (June 1 to June 30 in a non-leap year). We refer to this annual metric as ‘June maximum temperature’. We calculated the mean June maximum temperature across years at each station as well as year-specific values for temperature at each station to evaluate the effect of temperature on morphological variation across space and time, respectively. Species-wide mean temperature values were calculated by taking the mean June maximum temperature across all stations for each species.

### Derivation of morphological indices

Two morphological indices were derived from data collected on body mass and wing length for each bird. The Size Index (*SI*) corresponds to the overall size of an individual, while the Wing Index (*WI*) corresponds to the relative (to body mass) wing length, or “wingyness”, of each individual. These indices were derived using the expected power law (*57, 58*) relationship between these two traits,

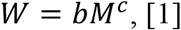

where *W* is wing length, *M* is body mass, *b* is a scalar, and *c* is the scaling exponent (Fig. 1B, 1C, S2), denoting how rapidly wing length increases as a function of mass. This relationship is linearized when taking the log of both sides of the equation,

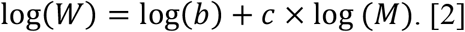

Using species-level mean values for both log(*W*) and log(*M*), we estimated the scaling exponent by applying a phylogenetic regression [to control for the effect of phylogenetic relatedness on parameter estimates (*59*)] using the ‘caper’ package (*60*) in R (*61*) to the linearized form of the power law relationship (Eq. 2). Species-level mean values were used because we were interested in understanding the general relationship between wing length and body mass. This represents the null expectation for how wing length covaries with mass. We estimated the scaling exponent for each of 100 phylogenetic trees for the species of interest obtained from BirdTree [(*62*) www.birdtree.org] to account for uncertainty in the phylogenetic relatedness of these species. The mean of the 100 estimates (mean = 0.333, standard deviation = 0.002) of the empirical relationship between wing length and body mass (i.e., the scaling exponent) was nearly identical to the theoretical expectation, given isometric scaling principles (where 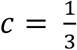; mass is expected to be proportional to volume, which scales as the cube of a linear dimension, such as wing length), and similar to estimates from other studies (*63, 64*) (Fig. S2).

For each species, measurements of body mass and wing length of individual bird captures were then reprojected onto new axes using a rotation matrix derived from the estimated scaling exponent (i.e., the rate at which wing length is expected to change with body mass). The rotation matrix was specified as,

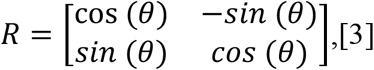

where *θ* is the amount (in radians) the data are to be rotated. We specified *θ* as the negative arc-tangent of *c* (as applying the arc-tangent function to the tangent of a triangle [the tangent being equivalent to the slope of a line] produces the angle in radians). For each species, we applied the rotation matrix to logged body mass (*LM*) and logged wing length (LW), to reproject the data onto new axes (Fig. S2),

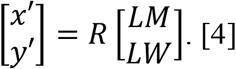

These reprojected data (*x*′ and *y*′) were standardized within species (i.e., centered and divided by the standard deviation) to create two relative indices (*SI* = size index; *WI* = wing index) that represent the overall size of the individual, and the degree to which wing length deviates from its expected value given the body mass of the individual, respectively,

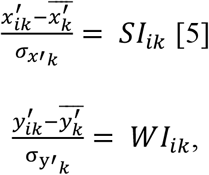

where 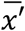 and 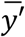 represent the mean and *σ*_*x*′_ and *σ*_*y*′_ represent the standard deviation of *x*′ and *y*′, respectively, for each species, *k*, and *i* represents each bird capture. This approach allowed us to account for the expected non-linear relationship among these traits when assessing spatiotemporal change and provides a means by which to assess morphological deviations from an expectation derived from empirical estimates rooted in scaling theory (*16*). *SI* values were closely correlated with logged mass (mean correlation coefficient across species = 0.99, range = 0.98 – 1). *WI* values showed a strong correlation to logged wing length (mean correlation coefficient across species = 0.75, range = 0.49 – 0.88), though not as strong as the relationship between *SI* and logged mass.

### Morphology as a function of time, latitude, and elevation

We used a hierarchical Bayesian approach to determine how *SI* and *WI* varied within species as a function of time, latitude, and elevation. We fit separate models for each index, that were identical in structure. In each case, the index (*y*_*IDX*_) for capture *i*, at banding station *j*, for species *k* was modeled as t-distributed, as a linear function of time,

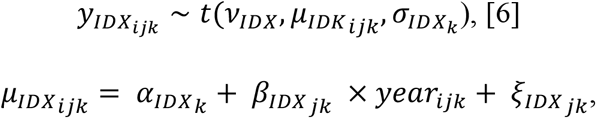

where *α*_*IDX*_ is the species-level intercept term, *β*_*IDX*_ is the effect of *year* on the response variable, *ξ*_*IDX*_ is the species-station intercept term, *σ*_*IDX*_ is the species-specific process error, *ν*_*IDX*_ represents the degrees of freedom, controlling the normality of the distribution (resulting in a Cauchy distribution when *ν*_*IDX*_ = 1 and approaching a normal distribution as *ν*_*IDX*_ approaches infinity), and the *IDX* subscript denotes the association of that parameter with this model (to help distinguish these parameters from those in other models). The degrees of freedom parameter of the t-distribution allows for additional flexibility (compared with the normal distribution) in modeling the structure of the residuals [for instance when there are ‘extreme observations’ (*65*)]. Parameter *α*_*IDX*_ was modeled as normally distributed,

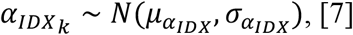

where 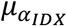 and 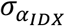 represent the mean and standard deviation of *α*_*IDX*_ across all species, respectively. Parameter *β*_*IDX*_ was modeled as normally distributed,

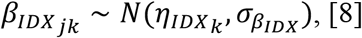

where *η*_*IDX*_ represents the mean effect of year on the response for each species, and 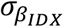 represents the process error. Parameter *σ*_*IDX*_ was modeled as half-normal (normal but with support only over positive values),

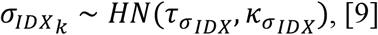

where 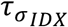 and 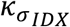 represent the mean and standard deviation of *σ*_*IDX*_, respectively. Process error was modeled hierarchically, as the degree to which these explanatory variables explain variation in the data may vary by species. Parameter *η*_*IDX*_ was modeled as normally distributed,

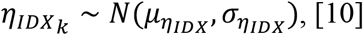

where 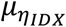 and 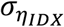 represent the mean and standard deviation of *η*_*IDX*_ across all species, respectively. The species-station intercept term, *ξ*_*IDX*_, was modeled as a linear function of latitude and elevation,

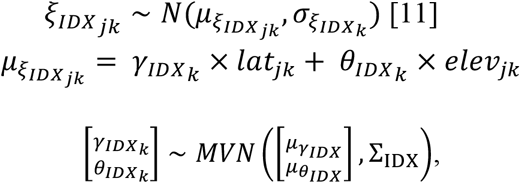

where *γ*_*IDX*_ is the species-specific effect of latitude (*lat*) on *ξ*_*IDX*_, *θ*_*IDX*_ is the species-specific effect of elevation (*elev*) on *ξ*_*IDX*_, and 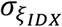 is the species-specific process error. Parameters *γ*_*IDX*_ and *θ*_*IDX*_ were modeled as multivariate normal, with means 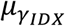 and 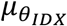, respectively, and covariance Σ_IDX_ (a 2 × 2 covariance matrix). Parameter 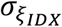 was modeled as half-normal

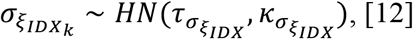

where 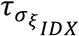 and 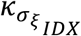 represent the mean and standard deviation of 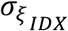, respectively. We fit all Bayesian models in this study using the R package ‘rstan’ (*66*) to interface with Stan (*67*) in R (*61*). R package ‘MCMCvis’ (*68*) was used to summarize, visualize, and manipulate all Bayesian model output. General data manipulation and processing was done using the ‘tidyverse’ family of R packages (*69*). For each model, we ran four chains for 8000 iterations each with a warmup of 4000 iterations. For all models, Rhat <= 1.01 and the number of effective samples was > 400 for all parameters. No models had divergent transitions (*67*). Weakly informative priors were given for all parameters. Graphical posterior predictive checks were used to check that data generated by the model were similar to the data used to fit the model (*70*). Data simulated from the posterior predictive distribution were similar to the observed data (Fig. S9).

For all model results in the main text, we present posterior mean estimates for parameters as well as the 89% credible intervals, following (*71*). The choice of 89% is arbitrary but serves to quantify parameter uncertainty while avoiding any suggestion that Bayesian credible intervals are analogous to tests of statistical significance (as might be assumed if using 95% cutoffs). For each parameter, we also present the probability that a given parameter is positive (calculated as the proportion of the posterior that is greater than 0) as *p*(> 0), or negative (the proportion of the posterior that is less than 0) as *p*(< 0). Scenarios in which *p*(> 0) or *p*(< 0) are near 0.5 indicate that a positive relationship is equally likely as a negative relationship.

To create species maps for Fig. 2C, 3C, 4B, we used range maps obtained from (*72*). Estimated effects of latitude and elevation were used to predict values for Size Index and Wing Index across the range of these species. We excluded all areas greater than 2000m of elevation (*73*) when deriving predictions for Size Index for red-eyed vireo (*Vireo olivaceus*) for Fig. 3C, to avoid making predictions outside the elevational range of this species in the Rocky Mountains.

### Body size as a function of temporal variation in temperature

To quantify how intraspecific variation in size across time is influenced by temperature, we modeled *SI* as a function *MT* (June maximum temperature at each station). The response variable (*y*_*TVT*_) for capture *i*, banding station *j*, and species *k* was modeled as t distributed, as a function of *MT*,

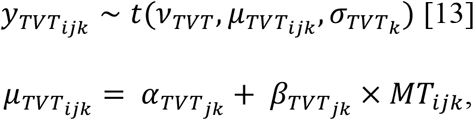

where *α*_*TVT*_ is the species-station-specific intercept term, *β*_*TVT*_ is the species-station-specific effect of temperature on the response variable, *σ*_*TVT*_ is the species-specific process error, *ν*_*TVT*_ represents the degrees of freedom, and the *TVT* subscript denotes the association of that parameter with this model. Parameter *α*_*TVT*_ was modeled normally distributed, as a function of *MST* (deviations of June maximum temperature from species-specific range-wide temperature at each station),

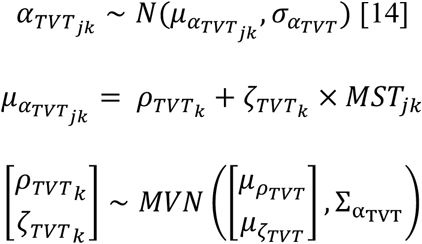

where *ρ*_*TVT*_ is the species-specific intercept term, *ζ*_*TVT*_ is the species-specific effects of MST on *α*_*TVT*_, and 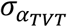 represents the process error. Parameters *ρ*_*TVT*_ and *ζ*_*TVT*_ were modeled as multivariate normal, with means 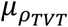 and 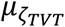, respectively and covariance 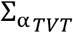 (a 2 × 2 covariance matrix). Parameter *β*_*TVT*_ was similarly modeled as a function of *MST*.

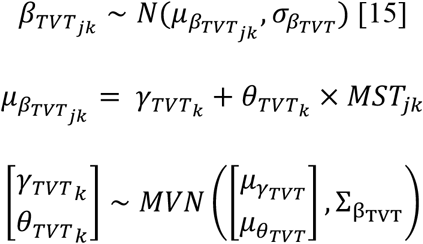

Both the intercept (*α*_*TVT*_) and slope (*β*_*TVT*_) at each species-station were modeled as a function of mean station temperature, because both the overall size and the effect of temporal variation in temperature might be expected to vary across this gradient. Parameter *σ*_*TVT*_ was modeled as half-normal,

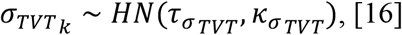

where 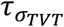 and 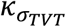 represent the mean and standard deviation of *σ*_*TVT*_, respectively. We fit three identical versions of this model, using temperature data in the year that the morphological data were collected (lag-0), as well as temperature one-(lag-1) and two-years (lag-2) prior to data collection, to explore the effect of temperature on morphology (i.e., the effect of temperature in year t, t − 1, and t − 2 on morphology in year t) during the potential hatching summer and subsequent summers and to account for the uncertainty and variability in the ages of these birds (all of which were known to be adults). For each model, we ran four chains for 6000 iterations each with a warmup of 3000 iterations.

### Body size as a function of spatial variation in temperature

To quantify how intraspecific variation in size across space is influenced by temperature, we modeled *SI* as a function of *MT* (mean June maximum temperature at each station across all years). The response variable (*y*_*SVT*_) for capture *i*, banding station *j*, and species *k* was modeled as t-distributed,

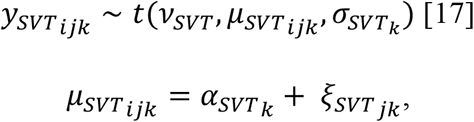

where *α*_*SVT*_ is the species-specific intercept term, *ξ*_*SVT*_ is the species-station-specific intercept, *σ*_*SVT*_ is the species-specific processes error, *ν*_*SVT*_ represents the degrees of freedom, and *SVT* denotes the association of each parameter with this model. Parameter *α*_*SVT*_ was modeled as normally distributed,

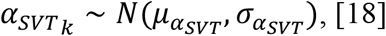

where *μ*_*α*_ and *σ*_*α*_ represent the mean and standard deviation of *α*_*SVT*_, respectively. Parameter *ξ*_*SVT*_ was modeled as normally distributed, as a function of *MT*,

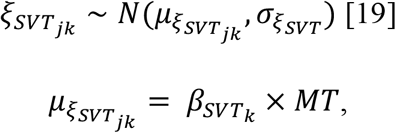

where 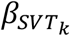 is the species-specific effect of *MT*, and 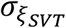 is the process error. Parameter *β*_*SVT*_ was modeled as normally distributed,

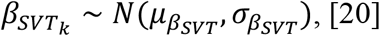

where 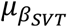 and 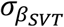 represent the mean and standard deviation of *β*_*SVT*_, respectively. We ran four chains for this model for 8000 iterations each with a warmup of 4000 iterations.

To assess how responses to temperature varied across species, we modeled the species-specific effect of spatial variation of temperature on *SI* (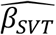; the posterior mean of *β*_*SVT*_ [Eq. 19], derived from the above model) and associated uncertainty as a function of *ST* (mean cross-station temperature within each species’ range). Parameter 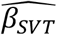 was modeled as normally distributed, with mean *π*_*SR*_ and standard deviation 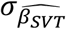 (the posterior standard deviation of *β*_*SVT*_ [Eq. 19], derived from the above model),

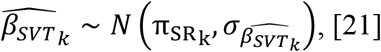

where SR denotes the association of each parameter with this model. In this way, the uncertainty in the species-specific estimates of the spatial temperature effect is propagated through these analyses. Parameter π_SVT_ was modeled as multivariate normal, as a linear function of *ST*, in a manner that accounts for the phylogenetic non-independence between species [following (*74, 75*)],

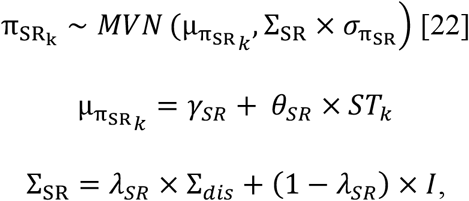

where *γ*_*SR*_ is the intercept term, *θ*_*SR*_ is the effect of *ST* on the response variable, and 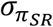 is the process error. Parameter Σ_*dis*_ is a phylogenetic covariance matrix, standardized such that the diagonal elements have a value of 1. The off-diagonal elements of Σ_*dis*_ describe the pair-wise phylogenetic distances between the 105 species included in this study. The phylogenetic covariance matrix was calculated from a consensus phylogenetic tree [calculated using the ‘phytools’ package (*76*) in R] based on 100 trees for the species of interest obtained from BirdTree [(*62*) www.birdtree.org]. Parameter *λ*_*SR*_ is Pagel’s lambda (*77*), which represents the degree to which phylogenetic relatedness contributes to variation in *π*_*SR*_, where values near 0 (the lower bound of the parameter) indicate low phylogenetic signal and values near 1 (the upper bound of the parameter) correspond to variation following a Brownian motion model of evolution (*75*), and *I* is an identity matrix. We ran this model for 1000 iterations with a warmup of 500 iterations.

### Back-transformation of effect sizes to trait space

Steps outlined by Eqs. 1-5 were implemented in reverse, to calculate the response of absolute morphological measurements (body mass and wing length) to variation over time, latitude, and elevation, using posterior estimates for the effects of these predictors on *SI* and *WI*. That is, for each species the effect sizes (i.e., posterior estimates) of these covariates on *SI* and *WI* were multiplied by the standard deviation of *x*′ and *y*′ (*σ*_*x*_′ and *σ*_*y*_′, respectively),

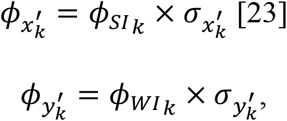

where *ϕ*_*SI*_ and *ϕ*_*WI*_ are the effect of a given covariate on *SI* and *WI*, respectively, for each species (*k*), and *ϕ*_*x*′_ and *ϕ*_*y*′_ represent the unstandardized effects of the covariate for each species. Parameters *ϕ*_*x*′_ and *ϕ*_*y*′_ were then rotated using the transpose of R (Eq. 3),

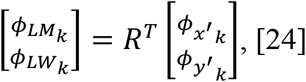

where *ϕ*_*LM*_ and *ϕ*_*LW*_ represent the effect of a given covariate on the logged absolute morphological metrics, *LM* (logged mass) and *LW* (logged wing length), for each species. This transformation has the effect of rotating data in the opposite direction of the rotation performed in Eq. 4. Since *ϕ*_*LM*_ and *ϕ*_*LW*_ represent an effect size in log space, when exponentiated, these metrics represent the multiplicative change in (unlogged) mass and wing length for each one-unit change in a given covariate. Subtracting one from this value and multiplying by 100 gives the percent change in that metric. To determine the percent change in mass (*ω*_*M*_) and wing length (*ω*_*W*_) over the temporal, latitudinal, and elevational range at which data were collected for each species, we exponentiated the product of *ϕ*_*LM*_ and *L* (for mass) and the product of *ϕ*_*LW*_ and *L* (for wing length), subtracted one, and multiplied by 100,

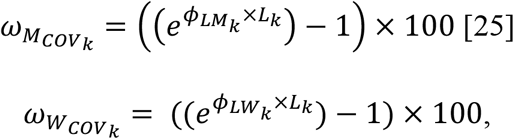

where *L* represents the total number of covariate units (i.e., 30 years, the latitudinal range in degrees for a given species, or the elevational range in meters for a given species), and *COV* represents time (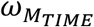 or 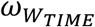), latitude (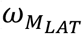 or 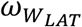), or elevation (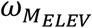 or 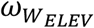). This was done at each iteration of the posterior for the estimated effect of year (*β*_*IDX*_; Eq. 6), latitude (*γ*_*IDX*_; Eq. 11), and elevation (*θ*_*IDX*_; Eq. 11), providing a posterior distribution for 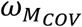 and 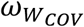. To calculate the cross-species mean percent change in mass and wing length, we calculated the mean of 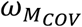 and 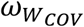 across all species at each posterior iteration, represented by 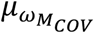 and 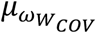, respectively,

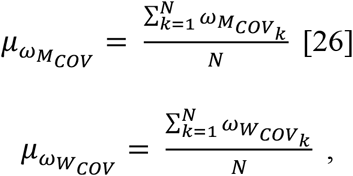

where N is the number of species.

### Rate of morphological change

To compare the observed rates of phenotypic change in this study to observed rates of evolutionary change in other taxa, we calculated change in logged mass in terms of *haldanes* (*h*),

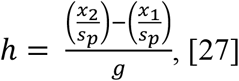

where *x*_2_ and *x*_1_ are the mean values for a morphological trait of interest at two time points, *s*_*p*_ is the standard deviation of the traits (pooled across time), and *g* is the number of generations that are likely to have occurred between the two time points (*78*). This measure, first proposed by (*79*), represents the magnitude of phenotypic change in standard deviations per generation.

For each species, we predicted logged mass at the beginning (*x*_1_) and end (*x*_2_) of the 30-year study period by subtracting and adding 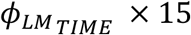 (where 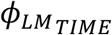 is from Eq. 24, representing change in logged mass per year), respectively, from mean logged mass. We calculated the within-population standard deviation across all years at each station and took the mean value of this standard deviation across stations (*s*_*p*_) for each species. We used information on generation length from (*80*) to calculate the number of generations (generation length / 30) for a particular species over this time period (*g*).

Prior work has suggested that rates of evolutionary change of |*h*| = 0.1 – 0.3 standard deviations per generation to be rapid (*81*), and that the maximal rate of phenotypic change that can be sustained indefinitely is approximately 0.1 phenotypic standard deviations per generation (*82*). For all species in this study, |*h*| < 0.1. Rates of phenotypic change were similar to those observed in other taxa undergoing anthropogenic disturbance (Fig. S10) (*83*).

## Acknowledgments

We thank MAPS station operators for collecting and sharing their data. Dani Kaschube provided critical data access, assisted by Rob Guralnick and Rafael LaFrance. We thank Christian Che-Castaldo for helpful discussions regarding the statistical modeling. Illustrations were provided by Lauren Helton.

## Author contributions

CY led formal analysis, and CY and MWT shared conceptualization and writing of the original draft. RBS and JFS facilitated data access. All authors contributed to review and editing of drafts.

## Data and Materials availability

All code used to produce analyses are freely available on Github (https://github.com/caseyyoungflesh/MAPS_morph_changes) and will be archived on Zenodo upon acceptance. Data from the Monitoring Avian Productivity and Survivorship (MAPS) program are curated and managed by The Institute for Bird Populations and were queried from the MAPS database on 2019-10-16. MAPS data will be archived on Dryad upon acceptance.

## Supplementary Materials

Figs. S1 to S10 Tables S1 to S4

## REFERENCES

1. D. Schluter, The evolution of finch communities on islands and continents: Kenya vs. Galapagos. Ecol. Monogr. 58, 229–249 (1988).

2. J. L. Gardner, A. Peters, M. R. Kearney, L. Joseph, R. Heinsohn, Declining body size: a third universal response to warming? Trends Ecol. Evol. 26, 285–291 (2011).

3. C. Bergmann, Über die Verhältnisse der wärmeökonomie der Thiere zu ihrer grösse. Gött. Stud. 3, 595–708 (1847).

4. K. Riemer, R. P. Guralnick, E. P. White, No general relationship between mass and temperature in endothermic species. eLife. 7, e27166 (2018).

5. D. M. Dehling, P. Jordano, H. M. Schaefer, K. Böhning-Gaese, M. Schleuning, Morphology predicts species’ functional roles and their degree of specialization in plant– frugivore interactions. Proc. R. Soc. B Biol. Sci. 283, 20152444 (2016).

6. P. R. Grant, Inheritance of size and shape in a population of Darwin’s finches, Geospiza conirostris. Proc. R. Soc. Lond. B Biol. Sci. 220, 219–236 (1983).

7. S. Des Roches, D. M. Post, N. E. Turley, J. K. Bailey, A. P. Hendry, M. T. Kinnison, J. A. Schweitzer, E. P. Palkovacs, The ecological importance of intraspecific variation. Nat. Ecol. Evol. 2, 57–64 (2018).

8. J. A. Allen, The influence of physical conditions in the genesis of species. Radic. Rev. 1, 108–140 (1877).

9. D. L. Altshuler, R. Dudley, The physiology and biomechanics of avian flight at high altitude. Integr. Comp. Biol. 46, 62–71 (2006).

10. C. Teplitsky, V. Millien, Climate warming and Bergmann’s rule through time: is there any evidence? Evol. Appl. 7, 156–168 (2014).

11. Y. Yom-Tov, S. Yom-Tov, J. Wright, C. J. R. Thorne, R. Du Feu, Recent changes in body weight and wing length among some British passerine birds. Oikos. 112, 91–101 (2006).

12. J. Van Buskirk, R. S. Mulvihill, R. C. Leberman, Declining body sizes in North American birds associated with climate change. Oikos. 119, 1047–1055 (2010).

13. B. C. Weeks, D. E. Willard, M. Zimova, A. A. Ellis, M. L. Witynski, M. Hennen, B. M. Winger, Shared morphological consequences of global warming in North American migratory birds. Ecol. Lett. 23, 316–325 (2020).

14. K. V. Rosenberg, A. M. Dokter, P. J. Blancher, J. R. Sauer, A. C. Smith, P. A. Smith, J. C. Stanton, A. Panjabi, L. Helft, M. Parr, P. P. Marra, Decline of the North American avifauna. Science. 366, 120–124 (2019).

15. D. F. DeSante, J. F. Saracco, D. R. O’Grady, K. M. Burton, B. L. Walker, Methodological considerations of the Monitoring Avian Productivity and Survivorship (MAPS) program. Stud. Avian Biol., 28–45 (2004).

16. G. B. West, J. H. Brown, B. J. Enquist, A general model for the origin of allometric scaling laws in biology. Science. 276, 122–126 (1997).

17. M. A. Ballinger, M. W. Nachman, The contribution of genetic and environmental effects to Bergmann’s rule and Allen’s rule in house mice. bioRxiv (2021).

18. S. C. Andrew, L. L. Hurley, M. M. Mariette, S. C. Griffith, Higher temperatures during development reduce body size in the zebra finch in the laboratory and in the wild. J. Evol. Biol. 30, 2156–2164 (2017).

19. A. M. Siepielski, M. B. Morrissey, S. M. Carlson, C. D. Francis, J. G. Kingsolver, K. D. Whitney, L. E. B. Kruuk, No evidence that warmer temperatures are associated with selection for smaller body sizes. Proc. R. Soc. B Biol. Sci. 286, 20191332 (2019).

20. E. A. Riddell, K. J. Iknayan, B. O. Wolf, B. Sinervo, S. R. Beissinger, Cooling requirements fueled the collapse of a desert bird community from climate change. Proc. Natl. Acad. Sci. 116, 21609–21615 (2019).

21. G. T. Pecl, M. B. Araújo, J. D. Bell, J. Blanchard, T. C. Bonebrake, I.-C. Chen, T. D. Clark, R. K. Colwell, F. Danielsen, B. Evengård, L. Falconi, S. Ferrier, S. Frusher, R. A. Garcia, R. B. Griffis, A. J. Hobday, C. Janion-Scheepers, M. A. Jarzyna, S. Jennings, J. Lenoir, H. Linnetved, V. Y. Martin, P. C. McCormack, J. McDonald, N. J. Mitchell, T. Mustonen, J. M. Pandolfi, N. Pettorelli, E. Popova, S. A. Robinson, B. R. Scheffers, J. D. Shaw, C. J. B. Sorte, J. M. Strugnell, J. M. Sunday, M.-N. Tuanmu, A. Vergés, C. Villanueva, T. Wernberg, E. Wapstra, S. E. Williams, Biodiversity redistribution under climate change: Impacts on ecosystems and human well-being. Science. 355, eaai9214 (2017).

22. D. J. Futuyma, Evolutionary constraint and ecological consequences. Evolution. 64, 1865–1884 (2010).

23. C. J. Murren, J. R. Auld, H. Callahan, C. K. Ghalambor, C. A. Handelsman, M. A. Heskel, J. G. Kingsolver, H. J. Maclean, J. Masel, H. Maughan, D. W. Pfennig, R. A. Relyea, S. Seiter, E. Snell-Rood, U. K. Steiner, C. D. Schlichting, Constraints on the evolution of phenotypic plasticity: limits and costs of phenotype and plasticity. Heredity. 115, 293–301 (2015).

24. M. W. Baldwin, H. Winkler, C. L. Organ, B. Helm, Wing pointedness associated with migratory distance in common-garden and comparative studies of stonechats (Saxicola torquata). J. Evol. Biol. 23, 1050–1063 (2010).

25. I. Newton, The migration ecology of birds (Elsevier, 2010).

26. S. M. Clegg, J. F. Kelly, M. Kimura, T. B. Smith, Combining genetic markers and stable isotopes to reveal population connectivity and migration patterns in a Neotropical migrant, Wilson’s warbler (Wilsonia pusilla). Mol. Ecol. 12, 819–830 (2003).

27. C. P. Bell, Leap-frog migration in the fox sparrow: Minimizing the cost of spring migration. The Condor. 99, 470–477 (1997).

28. S. Billerman, B. Keeney, P. Rodewald, T. Schulenberg, Birds of the World. Ithaca N. Y. Cornell Lab. Ornithol. (2020).

29. A. Desrochers, Morphological response of songbirds to 100 years of landscape change in North America. Ecology. 91, 1577–1582 (2010).

30. J. P. Swaddle, R. Lockwood, Morphological adaptations to predation risk in passerines. J. Avian Biol. 29, 172–176 (1998).

31. S. L. Chown, C. J. Klok, Altitudinal body size clines: latitudinal effects associated with changing seasonality. Ecography. 26, 445–455 (2003).

32. W. A. Boyle, Altitudinal bird migration in North America. The Auk. 134, 443–465 (2017).

33. A. R. Spence, M. W. Tingley, Body size and environment influence both intraspecific and interspecific variation in daily torpor use across hummingbirds. Funct. Ecol. 35, 870–883 (2021).

34. R. E. Moreau, Variation in the western Zosteropidae (Aves). Bull. Br. Mus. Nat. Hist. Zool. 4, 311–433 (1957).

35. T. H. Hamilton, The adaptive significances of intraspecific trends of variation in wing length and body size among bird species. Evolution. 15, 180–194 (1961).

36. I. D. Hodkinson, Terrestrial insects along elevation gradients: species and community responses to altitude. Biol. Rev. 80, 489–513 (2005).

37. P. Feinsinger, R. K. Colwell, J. Terborgh, S. B. Chaplin, Elevation and the morphology, flight energetics, and foraging ecology of tropical hummingbirds. Am. Nat. 113, 481–497 (1979).

38. J. W. Aldrich, Ecogeographical variation in size and proportions of song sparrows (Melospiza melodia). Ornithol. Monogr., 1–134 (1984).

39. Y. Sun, M. Li, G. Song, F. Lei, D. Li, Y. Wu, The role of climate factors in geographic variation in body mass and wing length in a passerine bird. Avian Res. 8, 1–9 (2017).

40. C. Sheard, M. H. C. Neate-Clegg, N. Alioravainen, S. E. I. Jones, C. Vincent, H. E. A. MacGregor, T. P. Bregman, S. Claramunt, J. A. Tobias, Ecological drivers of global gradients in avian dispersal inferred from wing morphology. Nat. Commun. 11 (2020).

41. S. Des Roches, L. H. Pendleton, B. Shapiro, E. P. Palkovacs, Conserving intraspecific variation for nature’s contributions to people. Nat. Ecol. Evol. 5, 574–582 (2021).

42. E. A. Riddell, K. J. Iknayan, B. O. Wolf, B. Sinervo, S. R. Beissinger, Cooling requirements fueled the collapse of a desert bird community from climate change. Proc. Natl. Acad. Sci. 116, 21609–21615 (2019).

43. A. E. McKechnie, B. O. Wolf, Climate change increases the likelihood of catastrophic avian mortality events during extreme heat waves. Biol. Lett. 6, 253–256 (2010).

44. S. R. Conradie, S. M. Woodborne, S. J. Cunningham, A. E. McKechnie, Chronic, sublethal effects of high temperatures will cause severe declines in southern African arid-zone birds during the 21st century. Proc. Natl. Acad. Sci. 116, 14065–14070 (2019).

45. V. Radchuk, T. Reed, C. Teplitsky, M. van de Pol, A. Charmantier, C. Hassall, P. Adamík, F. Adriaensen, M. P. Ahola, P. Arcese, J. Miguel Avilés, J. Balbontin, K. S. Berg, A. Borras, S. Burthe, J. Clobert, N. Dehnhard, F. de Lope, A. A. Dhondt, N. J. Dingemanse, H. Doi, T. Eeva, J. Fickel, I. Filella, F. Fossøy, A. E. Goodenough, S. J. G. Hall, B. Hansson, M. Harris, D. Hasselquist, T. Hickler, J. Joshi, H. Kharouba, J. G. Martínez, J.-B. Mihoub, J. A. Mills, M. Molina-Morales, A. Moksnes, A. Ozgul, D. Parejo, P. Pilard, M. Poisbleau, F. Rousset, M.-O. Rödel, D. Scott, J. C. Senar, C. Stefanescu, B. G. Stokke, T. Kusano, M. Tarka, C. E. Tarwater, K. Thonicke, J. Thorley, A. Wilting, P. Tryjanowski, J. Merilä, B. C. Sheldon, A. Pape Møller, E. Matthysen, F. Janzen, F. S. Dobson, M. E. Visser, S. R. Beissinger, A. Courtiol, S. Kramer-Schadt, Adaptive responses of animals to climate change are most likely insufficient. Nat. Commun. 10 (2019).

46. E. A. Riddell, K. J. Iknayan, L. Hargrove, S. Tremor, J. L. Patton, R. Ramirez, B. O. Wolf, S. R. Beissinger, Exposure to climate change drives stability or collapse of desert mammal and bird communities. Science. 371, 633–636 (2021).

47. M. W. Tingley, W. B. Monahan, S. R. Beissinger, C. Moritz, Birds track their Grinnellian niche through a century of climate change. Proc. Natl. Acad. Sci. 106, 19637–19643 (2009).

48. C. Youngflesh, J. Socolar, B. R. Amaral, A. Arab, R. P. Guralnick, A. H. Hurlbert, R. LaFrance, S. J. Mayor, D. A. W. Miller, M. W. Tingley, Migratory strategy drives species-level variation in bird sensitivity to vegetation green-up. Nat. Ecol. Evol. 5, 987–994 (2021).

49. L. Blueweiss, H. Fox, V. Kudzma, D. Nakashima, R. Peters, S. Sams, Relationships between body size and some life history parameters. Oecologia. 37, 257–272 (1978).

50. M. Kleiber, Body size and metabolic rate. Physiol. Rev. 27, 511–541 (1947).

51. P. Yodzis, S. Innes, Body size and consumer-resource dynamics. Am. Nat. 139, 1151–1175 (1992).

52. R. O. Prum, Interspecific social dominance mimicry in birds: Social mimicry in birds. Zool. J. Linn. Soc. 172, 910–941 (2014).

53. P. Pyle, Identification guide to North American birds: a compendium of information on identifying, ageing, and sexing” near-passerines” and passerines in the hand (Slate Creek Press, 1997).

54. C. Leys, C. Ley, O. Klein, P. Bernard, L. Licata, Detecting outliers: Do not use standard deviation around the mean, use absolute deviation around the median. J. Exp. Soc. Psychol. 49, 764–766 (2013).

55. J. J. Danielson, D. B. Gesch, Global multi-resolution terrain elevation data 2010 (GMTED2010) (US Department of the Interior, US Geological Survey, 2011).

56. M. M. Thornton, R. Shrestha, Y. Wei, P. E. Thornton, S. Kao, B. E. Wilson, Daymet: Daily surface weather data on a 1-km grid for North America, version 4 (ORNL Distributed Active Archive Center, 2020).

57. C. H. Greenewalt, The flight of birds: the significant dimensions, their departure from the requirements for dimensional similarity, and the effect on flight aerodynamics of that departure. Trans. Am. Philos. Soc. 65, 1–67 (1975).

58. G. Longo, M. Montévil, Perspectives on Organisms: Biological Time, Symmetries, and Singularities (Springer, 2014).

59. P. H. Harvey, “Why and how phylogenetic relationships should be incorporated into studies of scaling” in Scaling in biology (Oxford University Press, 2000), p. 253–265.

60. D. Orme, R. Freckleton, G. Thomas, T. Petzoldt, S. Fritz, N. Isaac, W. Pearse, The caper package: comparative analysis of phylogenetics and evolution in R. R Package Version. 5, 1–36 (2013).

61. R Core Team, R: A language and environment for statistical computing (R Foundation for Statistical Computing, Vienna, Austria, 2021; https://www.R-project.org/).

62. W. Jetz, G. H. Thomas, J. B. Joy, K. Hartmann, A. O. Mooers, The global diversity of birds in space and time. Nature. 491, 444–448 (2012).

63. R. L. Nudds, G. W. Kaiser, G. J. Dyke, Scaling of avian primary feather length. PLoS ONE. 6, e15665 (2011).

64. R. Nudds, Wing-bone length allometry in birds. J. Avian Biol. 38, 515–519 (2007).

65. S. C. Anderson, T. A. Branch, A. B. Cooper, N. K. Dulvy, Black-swan events in animal populations. Proc. Natl. Acad. Sci. 114, 3252–3257 (2017).

66. Stan Development Team. 2018. Stan Modeling Language Users Guide and Reference Manual, Version 2.18.0. http://mc-stan.org.

67. B. Carpenter, A. Gelman, M. D. Hoffman, D. Lee, B. Goodrich, M. Betancourt, M. Brubaker, J. Guo, P. Li, A. Riddell, Stan : A probabilistic programming language. J. Stat. Softw. 76 (2017).

68. C. Youngflesh, MCMCvis: Tools to visualize, manipulate, and summarize MCMC output. J. Open Source Softw. 3, 640 (2018).

69. H. Wickham, M. Averick, J. Bryan, W. Chang, L. McGowan, R. François, G. Grolemund Hayes, L. Henry, J. Hester, M. Kuhn, T. Pedersen, E. Miller, S. Bache, K. Müller, J. Ooms, D. Robinson, D. Seidel, V. Spinu, K. Takahashi, D. Vaughan, C. Wilke, K. Woo, H. Yutani, Welcome to the Tidyverse. J. Open Source Softw. 4, 1686 (2019).

70. J. Gabry, D. Simpson, A. Vehtari, M. Betancourt, A. Gelman, Visualization in Bayesian workflow. J. R. Stat. Soc. Ser. A Stat. Soc. 182, 389–402 (2019).

71. R. McElreath, Statistical rethinking: A Bayesian course with examples in R and Stan (Chapman and Hall/CRC, 2018).

72. BirdLife International. 2019. Data zone. Available from http://www.birdlife.org/datazone/index.html.

73. S. Cramp, C. Perrins, Handbook of the birds of Europe, the Middle East and North Africa. The birds of the Western Palearctic. Crows to finches, vol. VIII. (Oxford University Press, 1994).

74. J. Che-Castaldo, C. Che-Castaldo, M. C. Neel, Predictability of demographic rates based on phylogeny and biological similarity. Conserv. Biol. 32, 1290–1300 (2018).

75. P. de Villemereuil, J. A. Wells, R. D. Edwards, S. P. Blomberg, Bayesian models for comparative analysis integrating phylogenetic uncertainty. BMC Evol. Biol. 12, 102 (2012).

76. L. J. Revell, phytools: an R package for phylogenetic comparative biology (and other things): phytools: R package. Methods Ecol. Evol. 3, 217–223 (2012).

77. M. Pagel, Inferring the historical patterns of biological evolution. Nature. 401, 877–884 (1999).

78. A. P. Hendry, M. T. Kinnison, Perspective: The pace of modern life: measuring rates of contemporary microevolution. Evolution. 53, 1637–1653 (1999).

79. P. Gingerich, Rates of evolution: effects of time and temporal scaling. Science. 222, 159–162 (1983).

80. J. P. Bird, R. Martin, H. R. Akçakaya, J. Gilroy, I. J. Burfield, S. Garnett, A. Symes, J. Taylor, Ç. H. Şekercioğlu, S. H. M. Butchart, Generation lengths of the world’s birds and their implications for extinction risk. Conserv. Biol. (2020).

81. P. D. Gingerich, Rates of evolution. Annu. Rev. Ecol. Evol. Syst. 40, 657–675 (2009).

82. R. Bürger, M. Lynch, Evolution and extinction in a changing environment: A quantitative-genetic analysis. Evolution. 49, 151–163 (1995).

83. A. P. Hendry, T. J. Farrugia, M. T. Kinnison, Human influences on rates of phenotypic change in wild animal populations. Mol. Ecol. 17, 20–29 (2008).

