## Supplementary materials for "Abiotic conditions shape spatial and temporal morphological variation in North American birds"

**This PDF file includes:**

Figs. S1 to S10

Tables S1 to S4

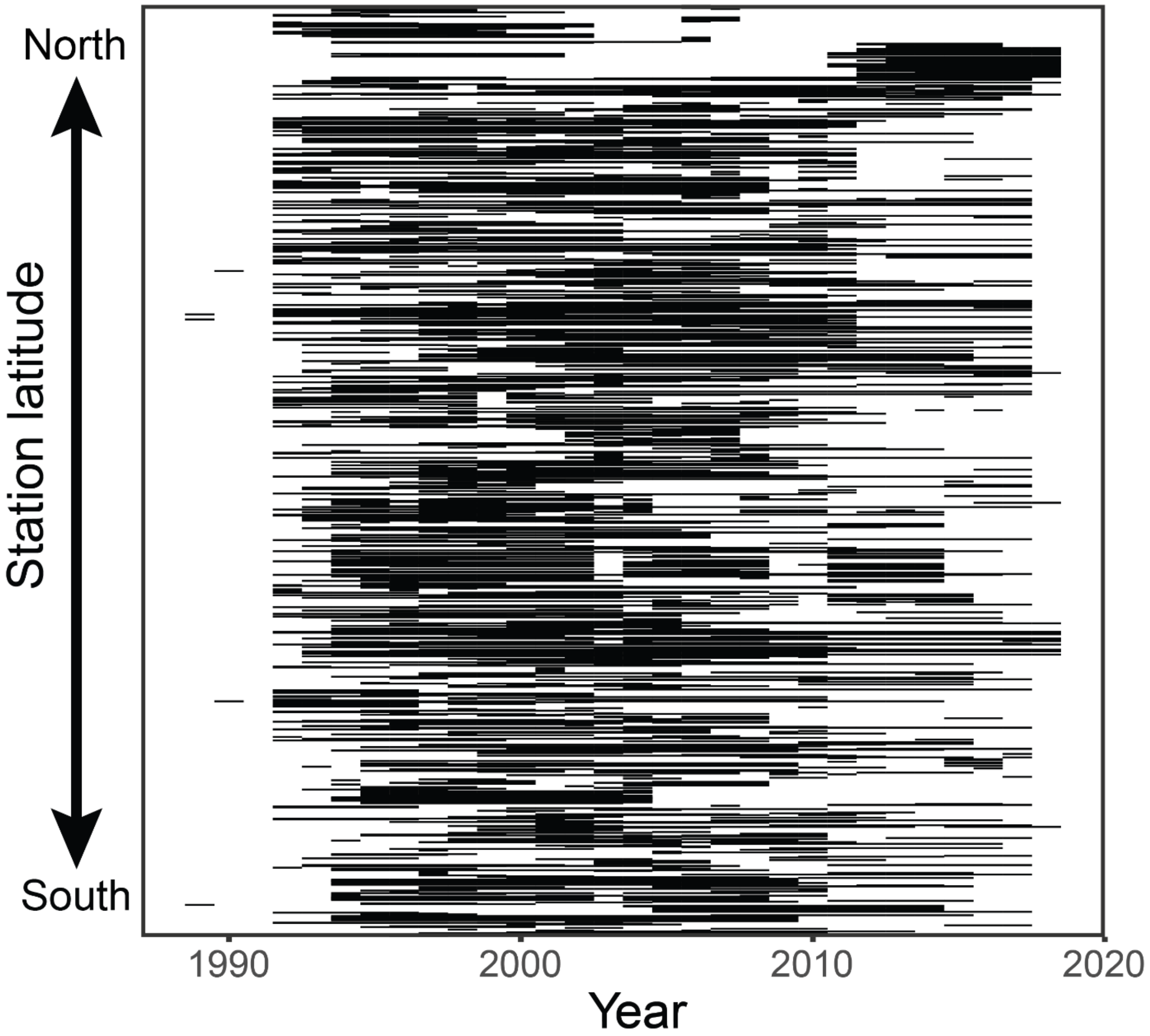


**Fig. S1. Morphological data availability over time**. Each horizontal line represents one of 1124 MAPS stations. Stations are ordered by latitude, from North (top) to South (bottom).

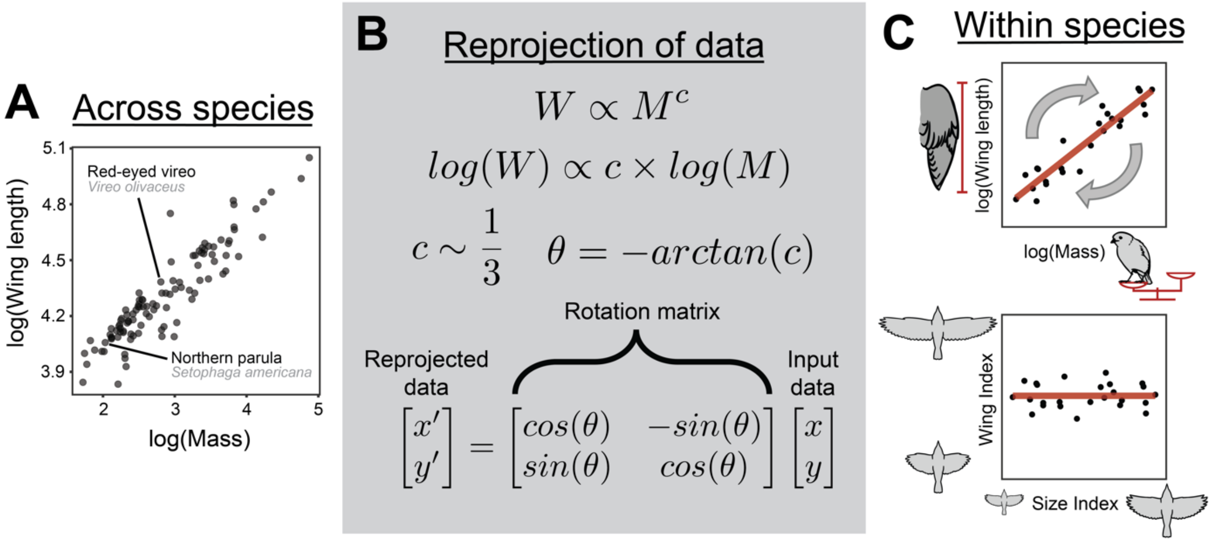


**Fig. S2. Derivation of morphological indices.** (**A**) Logged wing length as a function of logged mass for the 105 bird species considered in this study. Points represent mean values for each species. (**B**) The relationship between wing length (*W*) and mass (*M*) can be described by a power law, where *c* represents the scaling exponent. Logging both sides of the equation linearizes this model. Using a phylogenetic regression, *c* was estimated to be approximately 1/3 across species, as predicted by scaling theory. The negative arc tangent of this estimate (to convert the slope to radians) was used to create a rotation matrix. (**C**) For each species, the rotation matrix was used to reproject logged wing length and logged mass onto a new coordinate plane (top panel). Values for both the x and y axes were standardized to have a standard deviation of 1, to create a Size Index and Wing Index, representing the overall size of each individual bird and the degree to which wing length deviates from its expected value given the body mass of the individual, respectively (bottom panel).

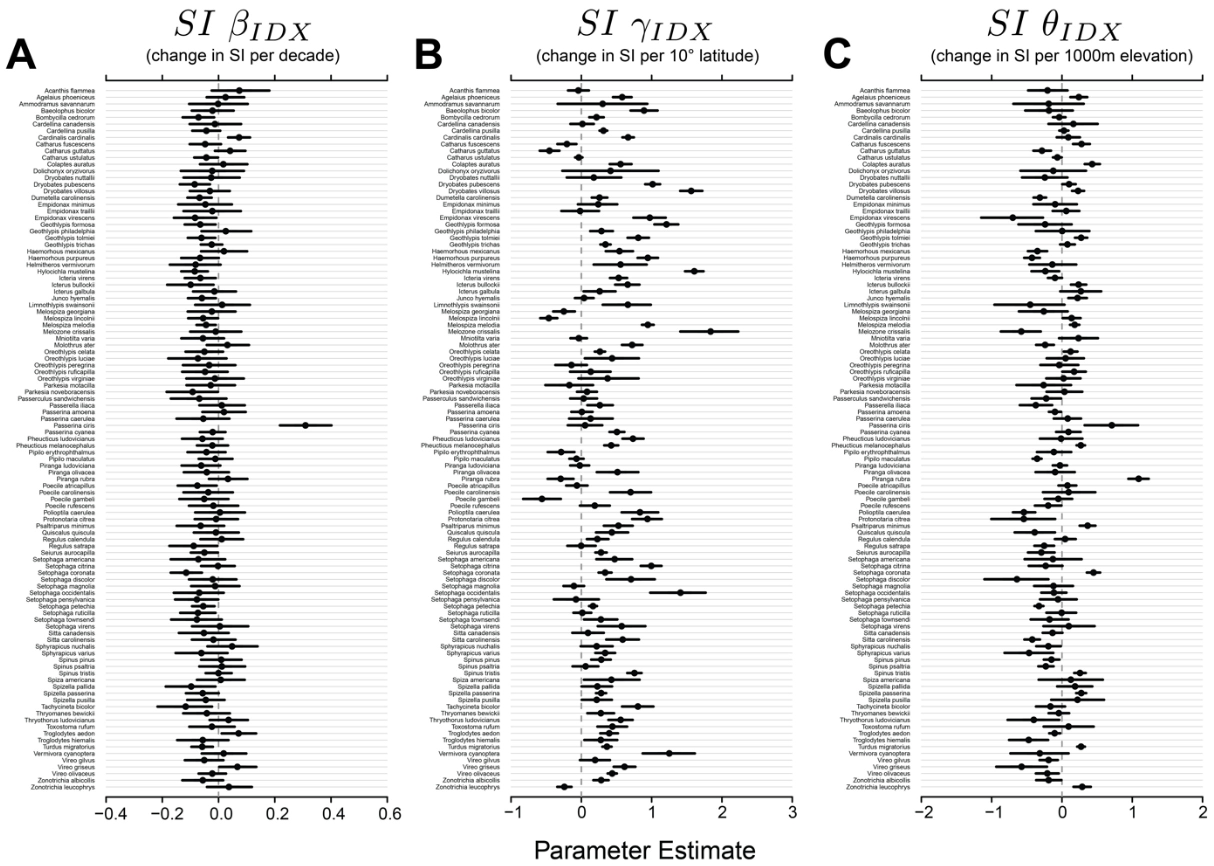


**Fig. S3. Posterior estimates for (A) $SI\ \beta_{IDX}$ (Eq. 6), (B) $SI\ \gamma_{IDX}$ (Eq. 11), and (C) $SI\ \theta_{IDX}$ (Eq. 11), denoting the change in Size Index for each species per 10 years, 10 degrees latitude, and 1000m elevation, respectively.** Points represent the posterior medians, while lines represent the 89% credible intervals. The dashed grey line represents zero in all cases.

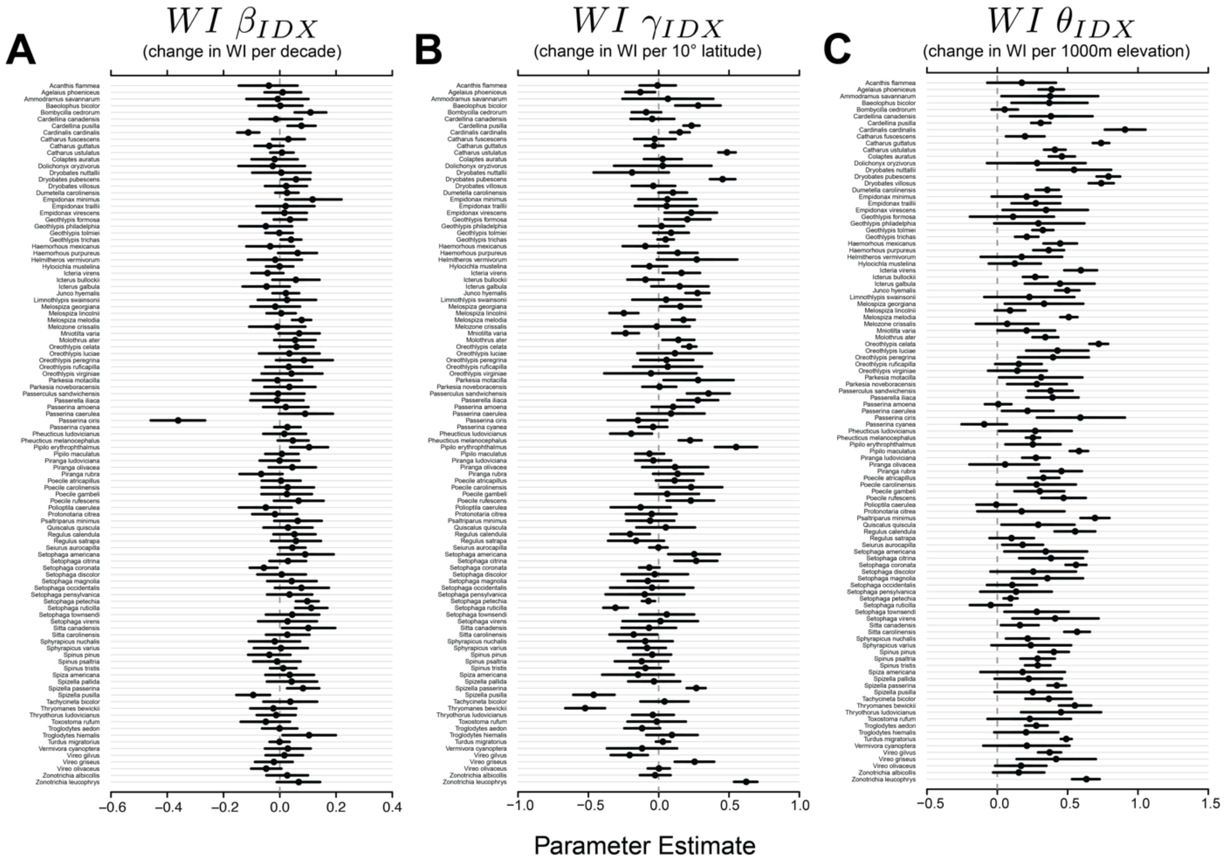


**Fig. S4. Posterior estimates for (A) $WI\ \beta_{IDX}$ (Eq. 6), (B) $WI\ \gamma_{IDX}$ (Eq. 11), and (C) $WI\ \theta_{IDX}$ (Eq. 11), denoting the change in Wing Index for each species per 10 years, 10 degrees latitude, and 1000m elevation, respectively.** Points represent the posterior medians, while lines represent the 89% credible intervals. The dashed grey line represents zero in all cases.

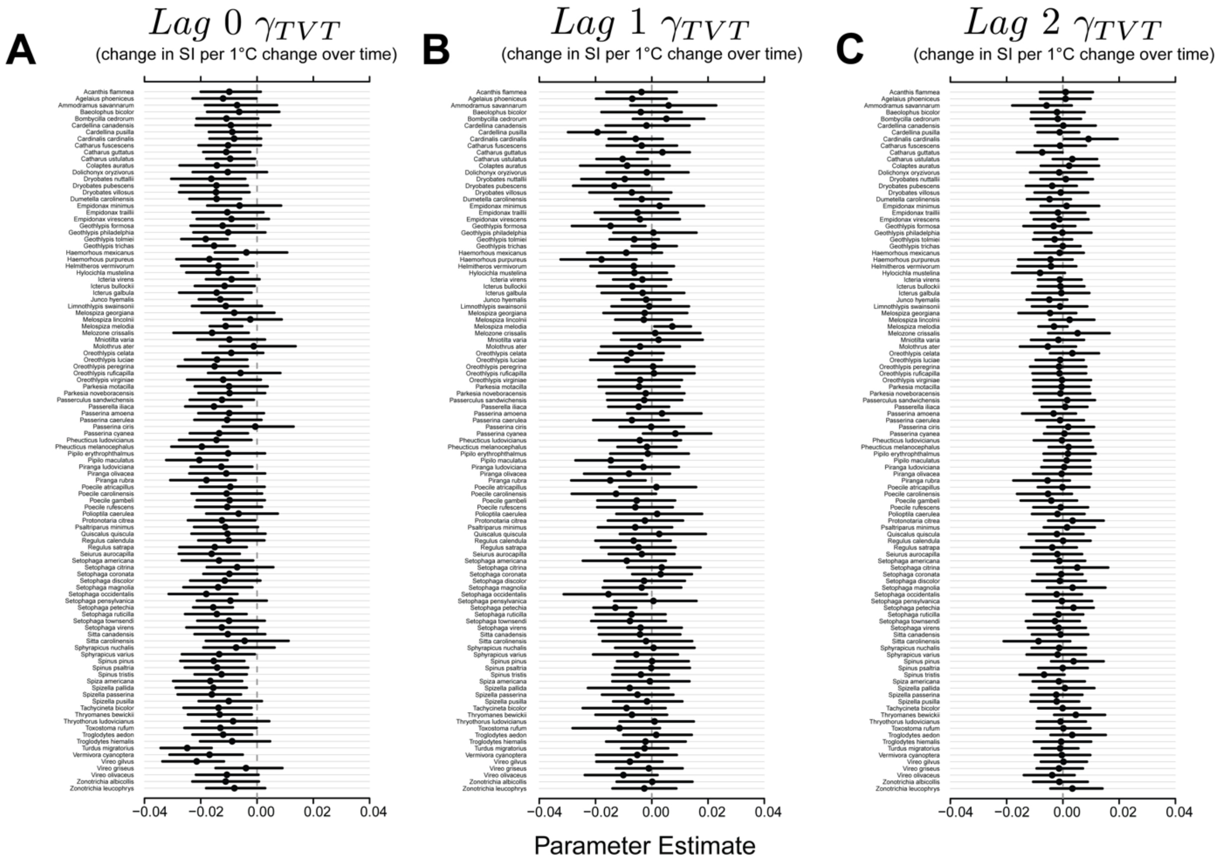


**Fig. S5. Posterior estimates for (A) $Lag\ 0\ \gamma_{TVT}$ (Eq. 15), (B) $Lag\ 1\ \gamma_{TVT}$ (Eq. 15), and (C) $Lag\ 2\ \gamma_{TVT}$ (Eq. 15), denoting the change in Size Index for each species per 1°C change in temperature at lag 0, lag 1, lag 2, respectively (i.e., the effect of change in temperature over time).** Points represent the posterior medians, while lines represent the 89% credible intervals. The dashed grey line represents zero in all cases.

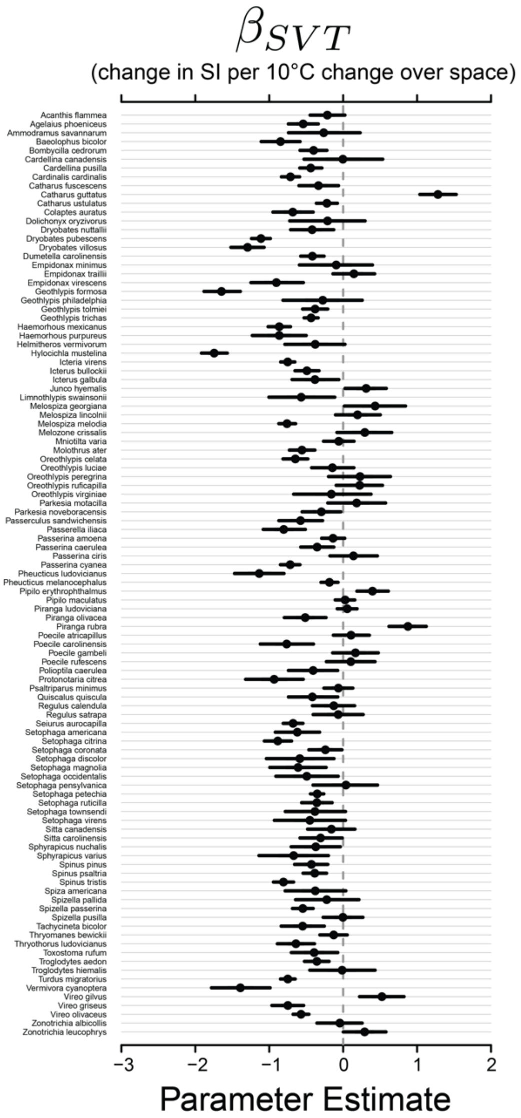


**Fig. S6. Posterior estimates for $\boldsymbol{\beta_{SVT}}$ (Eq. 19) denoting the change in Size Index for each species per 10°C change in mean station temperature (i.e., the effect of change in temperature over space).** Points represent the posterior medians, while lines represent the 89% credible intervals. The dashed grey line represents zero in all cases.

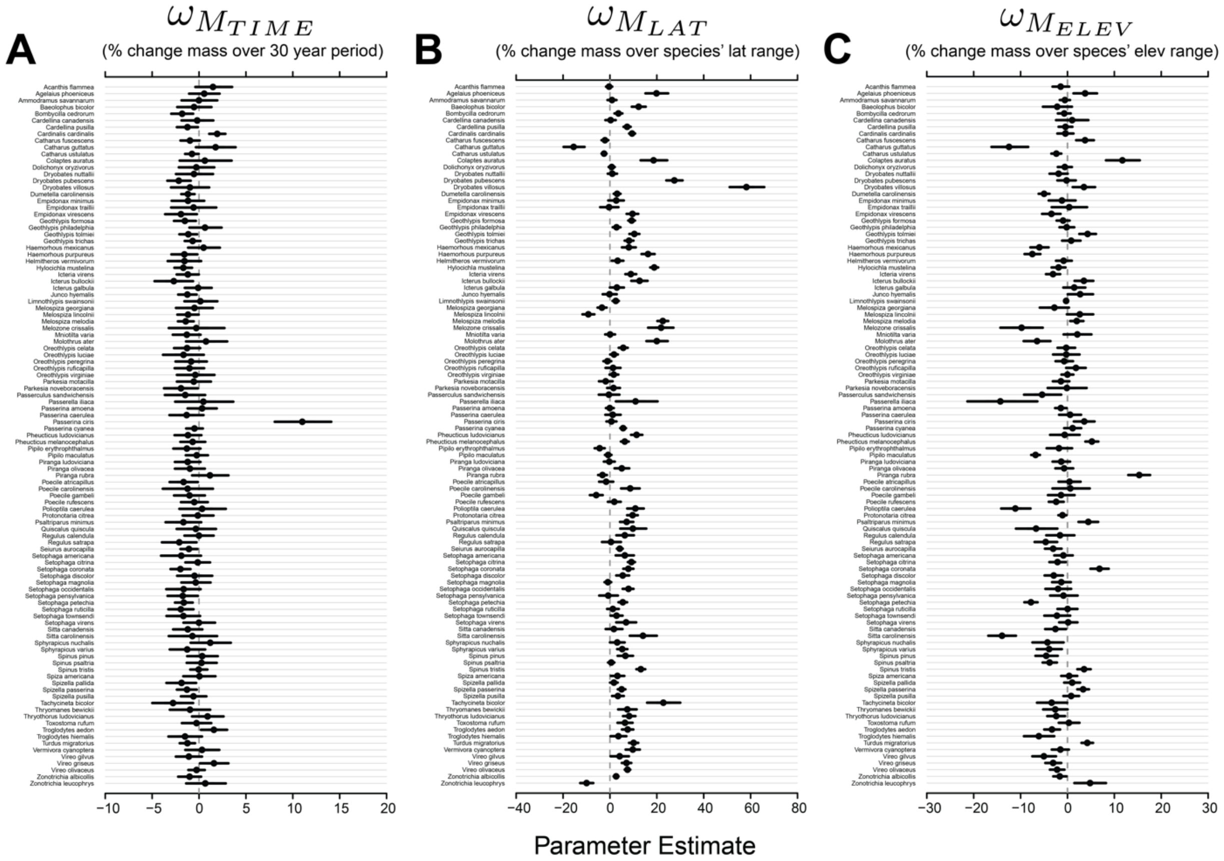


**Fig. S7. Posterior estimates for (A) $\omega_{M_{TIME}}$ (Eq. 25), (B) $\omega_{M_{LAT}}$ (Eq. 25), and (C) $\omega_{M_{ELEV}}$ (Eq. 25), denoting the percent change in mass for each species over the 30-year study period, the latitudinal range across which each species was sampled, and the elevational range across which each species was sampled, respectively.** Points represent the posterior medians, while lines represent the 89% credible intervals. The dashed grey line represents zero in all cases.

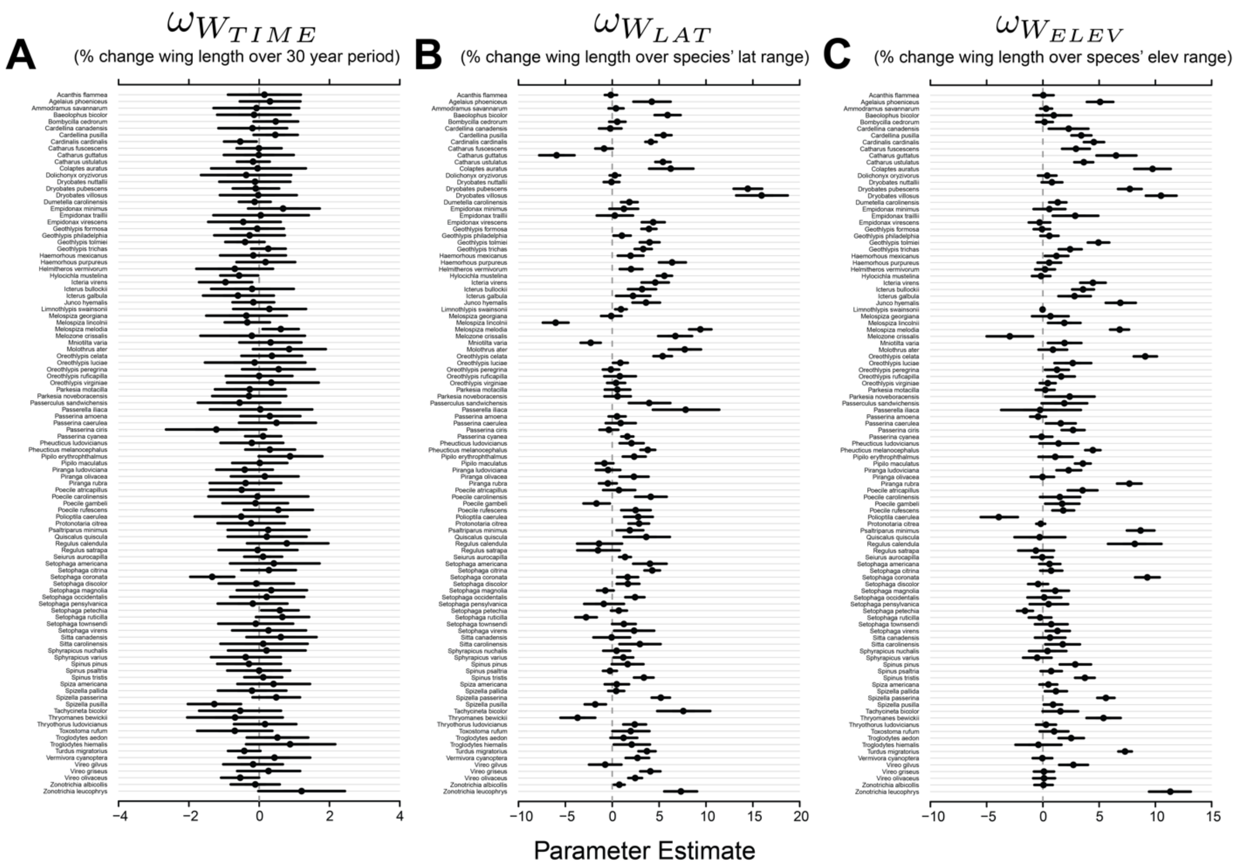


**Fig. S8. Posterior estimates for (A) $\omega_{W_{TIME}}$ (Eq. 25), (B) $\omega_{W_{LAT}}$ (Eq. 25), and (C) $\omega_{W_{ELEV}}$ (Eq. 25), denoting the percent change in wing length for each species over the 30-year study period, the latitudinal range across which each species was sampled, and the elevational range across which each species was sampled, respectively.** Points represent the posterior medians, while lines represent the 89% credible intervals. The dashed grey line represents zero in all cases.

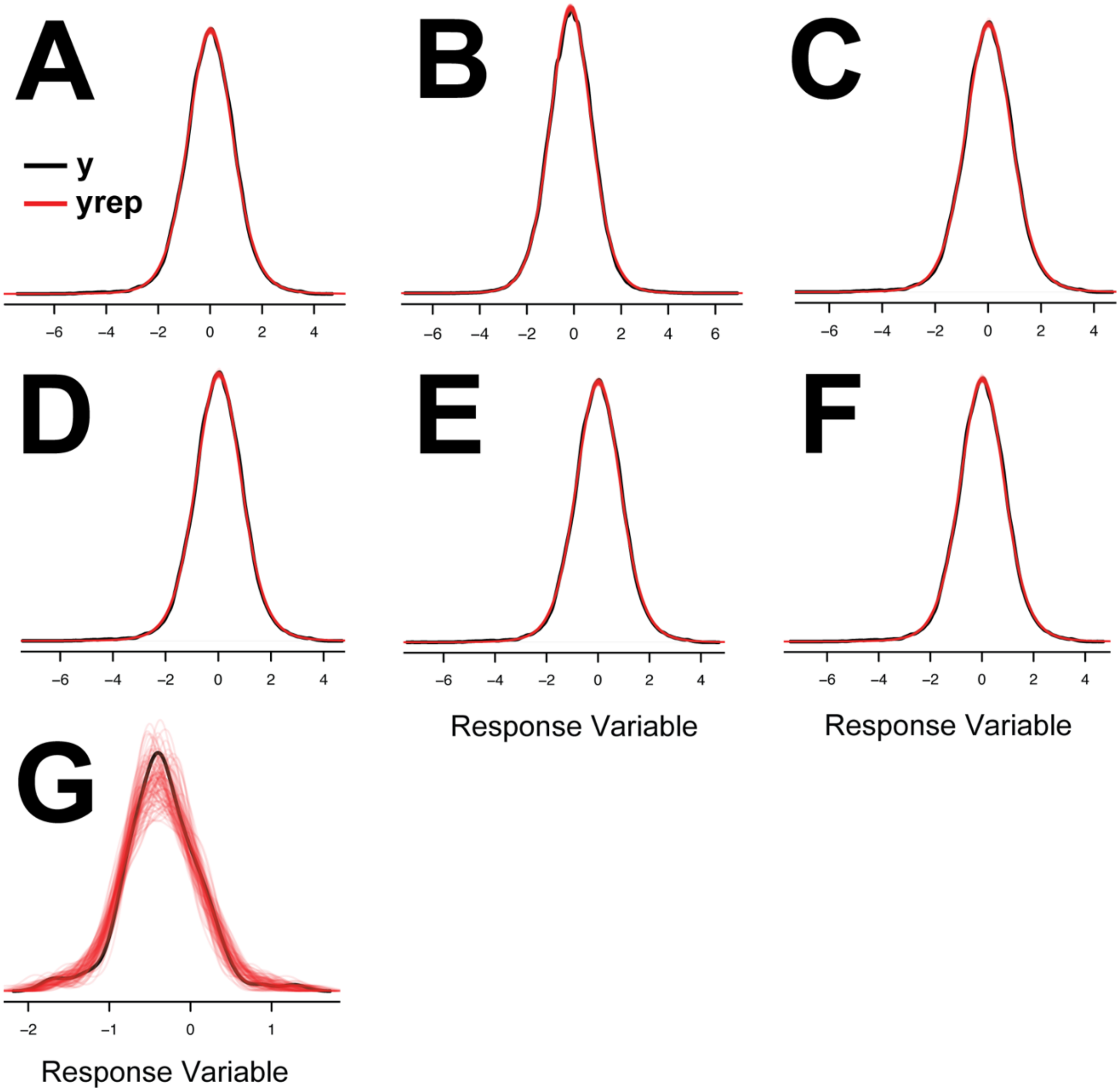


**Fig. S9. Density plots for observed response variable data (*y*) and response data simulated from the posterior predictive distribution (*yrep*).** These plots were used for graphical posterior predictive checks, to ensure that data simulated from the model were similar to the observed data for models examining (**A**) how Size Index varies in response to time, latitude, and elevation (Eqs. 6–12), (**B**) how Wing Index varies in response to time, latitude, and elevation (Eqs. 6–12), (**C**) how Size Index varies in response to temporal variation in temperature at lag 0 (Eqs. 13–16), (**D**) how Size Index varies in response to temporal variation in temperature at lag 1 (Eqs. 13–16),

(**E**) how Size Index varies in response to temporal variation in temperature at lag 2 (Eqs. 13–16), (**F**) how Size Index varies in response to spatial variation in temperature (Eqs. 17–20), and (**G**) how the effect of spatial variation in temperature on Size Index varies in response to the mean (range-wide) temperature experienced by that species (Eqs. 21–22). Curves in black are a representation of the density of all response data used to fit each model. Curves in red are a representation of the density of data simulated from the posterior predictive distribution. In some cases, black curves are obscured by the overlapping red curves. Each iteration of the posterior chain yields a simulated dataset. Here 100 datasets simulated from the posterior predictive distribution are displayed (100 separate red lines). The general similarities between the red lines and black lines demonstrate that the models simulate data similar to the observed data.

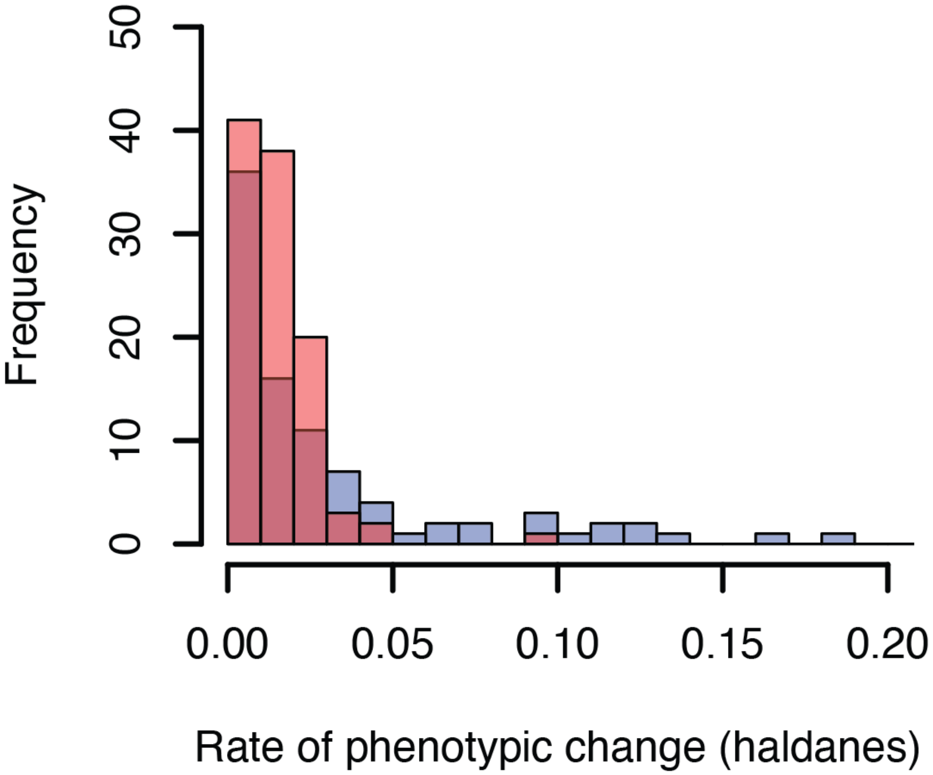


**Fig. S10. Absolute value of the estimate rate of change (represented in units of haldanes [standard deviations per generation]) for body mass ($|h|$) for focal species in this study (red) and for species and traits presented in (*79*) (blue)**. Traits considered by (*79*) varied by species and only species undergoing anthropogenic disturbance [as defined by (*79*)] were considered. The x-axis of the plot is truncated at 0.2 to facilitate visualization.

*Table S1*: For all species considered in this study, the scientific species name, English common name, family, order, number of captures (N), number of stations, latitudinal range of sampling, elevational range of sampling.

| Species | Common name | Family | Order | N | N stations | Lat range (degrees) | Elev range (m) |
|---|---|---|---|---|---|---|---|
| *Acanthis flammea* | Common Redpoll | Fringillidae | Passeriformes | 898 | 37 | 14.2 | 1032 |
| *Agelaius phoeniceus* | Red-winged Blackbird | Icteridae | Passeriformes | 2078 | 189 | 37.9 | 2511 |
| *Ammodramus savannarum* | Grasshopper Sparrow | Emberizidae | Passeriformes | 741 | 28 | 4.9 | 360 |
| *Baeolophus bicolor* | Tufted Titmouse | Paridae | Passeriformes | 666 | 188 | 16.7 | 1140 |
| *Bombycilla cedrorum* | Cedar Waxwing | Bombycillidae | Passeriformes | 3625 | 229 | 22.3 | 2156 |
| *Cardellina canadensis* | Canada Warbler | Parulidae | Passeriformes | 718 | 36 | 21.4 | 1581 |
| *Cardellina pusilla* | Wilson's Warbler | Parulidae | Passeriformes | 8543 | 184 | 33.7 | 2996 |
| *Cardinalis cardinalis* | Northern Cardinal | Cardinalidae | Passeriformes | 8580 | 422 | 19.6 | 1432 |
| *Catharus fuscescens* | Veery | Turdidae | Passeriformes | 3303 | 137 | 17.4 | 2439 |
| *Catharus guttatus* | Hermit Thrush | Turdidae | Passeriformes | 2274 | 192 | 29.3 | 2783 |
| *Catharus ustulatus* | Swainson's Thrush | Turdidae | Passeriformes | 10780 | 267 | 35.4 | 2996 |
| *Colaptes auratus* | Northern Flicker | Picidae | Piciformes | 382 | 178 | 28.7 | 2763 |
| *Dolichonyx oryzivorus* | Bobolink | Icteridae | Passeriformes | 458 | 20 | 3.3 | 577 |
| *Dryobates nuttallii* | Nuttall's Woodpecker | Picidae | Piciformes | 383 | 58 | 7.6 | 928 |
| *Dryobates pubescens* | Downy Woodpecker | Picidae | Piciformes | 2011 | 442 | 32.6 | 2601 |
| *Dryobates villosus* | Hairy Woodpecker | Picidae | Piciformes | 451 | 194 | 30.6 | 2783 |
| *Dumetella carolinensis* | Gray Catbird | Mimidae | Passeriformes | 12187 | 304 | 22.9 | 2447 |
| *Empidonax minimus* | Least Flycatcher | Tyrannidae | Passeriformes | 673 | 53 | 18.8 | 1525 |
| *Empidonax traillii* | Willow Flycatcher | Tyrannidae | Passeriformes | 401 | 57 | 18.3 | 2511 |
| *Empidonax virescens* | Acadian Flycatcher | Tyrannidae | Passeriformes | 871 | 122 | 13.0 | 612 |
| *Geothlypis formosa* | Kentucky Warbler | Parulidae | Passeriformes | 3536 | 142 | 10.6 | 515 |
| *Geothlypis philadelphia* | Mourning Warbler | Parulidae | Passeriformes | 584 | 38 | 15.5 | 648 |
| *Geothlypis tolmiei* | MacGillivray's Warbler | Parulidae | Passeriformes | 7306 | 185 | 19.5 | 2996 |
| *Geothlypis trichas* | Common Yellowthroat | Parulidae | Passeriformes | 13947 | 483 | 32.1 | 2511 |
| *Haemorhous mexicanus* | House Finch | Fringillidae | Passeriformes | 1687 | 128 | 20.6 | 2211 |
| *Haemorhous purpureus* | Purple Finch | Fringillidae | Passeriformes | 3075 | 120 | 23.6 | 2331 |
| *Helmitheros vermivorum* | Worm-eating Warbler | Parulidae | Passeriformes | 566 | 58 | 9.5 | 756 |
| *Hylocichla mustelina* | Wood Thrush | Turdidae | Passeriformes | 5251 | 260 | 16.1 | 1136 |
| *Icteria virens* | Yellow-breasted Chat | Parulidae | Passeriformes | 5031 | 214 | 24.0 | 2153 |
| *Icterus bullockii* | Bullock's Oriole | Icteridae | Passeriformes | 1328 | 105 | 21.0 | 2083 |
| *Icterus galbula* | Baltimore Oriole | Icteridae | Passeriformes | 729 | 96 | 20.7 | 1301 |
| *Junco hyemalis* | Dark-eyed Junco | Emberizidae | Passeriformes | 6385 | 237 | 33.7 | 2996 |

*Table S1*: For all species considered in this study, the scientific species name, English common name, family, order, number of captures (N), number of stations, latitudinal range of sampling, elevational range of sampling. *(continued)*

| Species | Common name | Family | Order | N | N stations | Lat range (degrees) | Elev range (m) |
|---|---|---|---|---|---|---|---|
| *Limnothlypis swainsonii* | Swainson's Warbler | Parulidae | Passeriformes | 675 | 31 | 6.4 | 102 |
| *Melospiza georgiana* | Swamp Sparrow | Emberizidae | Passeriformes | 835 | 60 | 16.7 | 1198 |
| *Melospiza lincolnii* | Lincoln's Sparrow | Emberizidae | Passeriformes | 3606 | 141 | 32.2 | 2981 |
| *Melospiza melodia* | Song Sparrow | Emberizidae | Passeriformes | 15640 | 466 | 26.9 | 2511 |
| *Melozone crissalis* | California Towhee | Emberizidae | Passeriformes | 519 | 67 | 9.9 | 1616 |
| *Mniotilta varia* | Black-and-white Warbler | Parulidae | Passeriformes | 1148 | 165 | 27.1 | 1593 |
| *Molothrus ater* | Brown-headed Cowbird | Icteridae | Passeriformes | 1051 | 226 | 26.8 | 2511 |
| *Oreothlypis celata* | Orange-crowned Warbler | Parulidae | Passeriformes | 3169 | 151 | 35.4 | 2763 |
| *Oreothlypis luciae* | Lucy's Warbler | Parulidae | Passeriformes | 656 | 23 | 5.9 | 1506 |
| *Oreothlypis peregrina* | Tennessee Warbler | Parulidae | Passeriformes | 1772 | 58 | 12.4 | 1161 |
| *Oreothlypis ruficapilla* | Nashville Warbler | Parulidae | Passeriformes | 1095 | 77 | 16.7 | 1877 |
| *Oreothlypis virginiae* | Virginia's Warbler | Parulidae | Passeriformes | 437 | 23 | 6.5 | 797 |
| *Parkesia motacilla* | Louisiana Waterthrush | Parulidae | Passeriformes | 536 | 78 | 12.3 | 650 |
| *Parkesia noveboracensis* | Northern Waterthrush | Parulidae | Passeriformes | 469 | 64 | 28.3 | 2440 |
| *Passerculus sandwichensis* | Savannah Sparrow | Emberizidae | Passeriformes | 787 | 44 | 31.4 | 2729 |
| *Passerella iliaca* | Fox Sparrow | Emberizidae | Passeriformes | 782 | 72 | 35.1 | 2996 |
| *Passerina amoena* | Lazuli Bunting | Emberizidae | Passeriformes | 1440 | 116 | 16.7 | 2153 |
| *Passerina caerulea* | Blue Grosbeak | Emberizidae | Passeriformes | 888 | 78 | 15.9 | 1853 |
| *Passerina ciris* | Painted Bunting | Emberizidae | Passeriformes | 1944 | 56 | 8.8 | 573 |
| *Passerina cyanea* | Indigo Bunting | Emberizidae | Passeriformes | 5458 | 288 | 16.6 | 1594 |
| *Pheucticus ludovicianus* | Rose-breasted Grosbeak | Cardinalidae | Passeriformes | 869 | 119 | 21.4 | 1579 |
| *Pheucticus melanocephalus* | Black-headed Grosbeak | Cardinalidae | Passeriformes | 3935 | 242 | 19.7 | 2763 |
| *Pipilo erythrophthalmus* | Eastern Towhee | Emberizidae | Passeriformes | 1238 | 222 | 16.5 | 1597 |
| *Pipilo maculatus* | Spotted Towhee | Emberizidae | Passeriformes | 3450 | 209 | 21.0 | 2341 |
| *Piranga ludoviciana* | Western Tanager | Cardinalidae | Passeriformes | 1867 | 167 | 25.9 | 2782 |
| *Piranga olivacea* | Scarlet Tanager | Cardinalidae | Passeriformes | 554 | 136 | 15.2 | 1033 |
| *Piranga rubra* | Summer Tanager | Cardinalidae | Passeriformes | 1145 | 154 | 11.3 | 1499 |
| *Poecile atricapillus* | Black-capped Chickadee | Paridae | Passeriformes | 1369 | 266 | 27.6 | 2511 |
| *Poecile carolinensis* | Carolina Chickadee | Paridae | Passeriformes | 419 | 135 | 12.2 | 1033 |
| *Poecile gambeli* | Mountain Chickadee | Paridae | Passeriformes | 606 | 89 | 17.0 | 2291 |
| *Poecile rufescens* | Chestnut-backed Chickadee | Paridae | Passeriformes | 597 | 81 | 23.2 | 1574 |
| *Polioptila caerulea* | Blue-gray Gnatcatcher | Polioptilidae | Passeriformes | 414 | 104 | 13.2 | 2379 |

*Table S1*: For all species considered in this study, the scientific species name, English common name, family, order, number of captures (N), number of stations, latitudinal range of sampling, elevational range of sampling. *(continued)*

| Species | Common name | Family | Order | N | N stations | Lat range (degrees) | Elev range (m) |
|---|---|---|---|---|---|---|---|
| *Protonotaria citrea* | Prothonotary Warbler | Parulidae | Passeriformes | 1531 | 71 | 13.8 | 268 |
| *Psaltriparus minimus* | Bushtit | Aegithalidae | Passeriformes | 1334 | 131 | 17.0 | 2341 |
| *Quiscalus quiscula* | Common Grackle | Icteridae | Passeriformes | 707 | 75 | 25.2 | 1885 |
| *Regulus calendula* | Ruby-crowned Kinglet | Reguliidae | Passeriformes | 787 | 77 | 33.5 | 2989 |
| *Regulus satrapa* | Golden-crowned Kinglet | Reguliidae | Passeriformes | 666 | 83 | 27.1 | 2443 |
| *Seiurus aurocapilla* | Ovenbird | Parulidae | Passeriformes | 4775 | 284 | 23.6 | 1539 |
| *Setophaga americana* | Northern Parula | Parulidae | Passeriformes | 391 | 72 | 19.2 | 654 |
| *Setophaga citrina* | Hooded Warbler | Parulidae | Passeriformes | 3130 | 122 | 12.8 | 1029 |
| *Setophaga coronata* | Yellow-rumped Warbler | Parulidae | Passeriformes | 3774 | 215 | 33.7 | 2996 |
| *Setophaga discolor* | Prairie Warbler | Parulidae | Passeriformes | 742 | 57 | 10.1 | 592 |
| *Setophaga magnolia* | Magnolia Warbler | Parulidae | Passeriformes | 938 | 64 | 16.1 | 1212 |
| *Setophaga occidentalis* | Hermit Warbler | Parulidae | Passeriformes | 617 | 46 | 7.8 | 2243 |
| *Setophaga pensylvanica* | Chestnut-sided Warbler | Parulidae | Passeriformes | 1101 | 61 | 17.5 | 1589 |
| *Setophaga petechia* | Yellow Warbler | Parulidae | Passeriformes | 12301 | 336 | 37.8 | 2996 |
| *Setophaga ruticilla* | American Redstart | Parulidae | Passeriformes | 3507 | 167 | 27.1 | 1397 |
| *Setophaga townsendi* | Townsend's Warbler | Parulidae | Passeriformes | 577 | 31 | 15.6 | 1556 |
| *Setophaga virens* | Black-throated Green Warbler | Parulidae | Passeriformes | 383 | 43 | 21.1 | 953 |
| *Sitta canadensis* | Red-breasted Nuthatch | Sittidae | Passeriformes | 398 | 127 | 26.9 | 2779 |
| *Sitta carolinensis* | White-breasted Nuthatch | Sittidae | Passeriformes | 498 | 161 | 19.6 | 2779 |
| *Sphyrapicus nuchalis* | Red-naped Sapsucker | Picidae | Piciformes | 530 | 69 | 16.8 | 2505 |
| *Sphyrapicus varius* | Yellow-bellied Sapsucker | Picidae | Piciformes | 417 | 81 | 21.2 | 1167 |
| *Spinus pinus* | Pine Siskin | Fringillidae | Passeriformes | 1406 | 102 | 29.7 | 2996 |
| *Spinus psaltria* | Lesser Goldfinch | Fringillidae | Passeriformes | 1391 | 96 | 11.0 | 2210 |
| *Spinus tristis* | American Goldfinch | Fringillidae | Passeriformes | 7665 | 333 | 24.5 | 2511 |
| *Spiza americana* | Dickcissel | Cardinalidae | Passeriformes | 947 | 46 | 10.5 | 559 |
| *Spizella pallida* | Clay-colored Sparrow | Emberizidae | Passeriformes | 1151 | 39 | 12.0 | 1104 |
| *Spizella passerina* | Chipping Sparrow | Emberizidae | Passeriformes | 2292 | 217 | 34.8 | 2783 |
| *Spizella pusilla* | Field Sparrow | Emberizidae | Passeriformes | 2690 | 142 | 16.5 | 646 |
| *Tachycineta bicolor* | Tree Swallow | Hirundinidae | Passeriformes | 594 | 62 | 33.6 | 2090 |
| *Thryomanes bewickii* | Bewick's Wren | Troglodytidae | Passeriformes | 1102 | 158 | 22.2 | 2098 |
| *Thryothorus ludovicianus* | Carolina Wren | Troglodytidae | Passeriformes | 1754 | 216 | 17.3 | 649 |

*Table S1*: For all species considered in this study, the scientific species name, English common name, family, order, number of captures (N), number of stations, latitudinal range of sampling, elevational range of sampling. *(continued)*

| Species | Common name | Family | Order | N | N stations | Lat range (degrees) | Elev range (m) |
|---|---|---|---|---|---|---|---|
| *Toxostoma rufum* | Brown Thrasher | Mimidae | Passeriformes | 554 | 109 | 22.0 | 953 |
| *Troglodytes aedon* | House Wren | Troglodytidae | Passeriformes | 2867 | 241 | 23.8 | 2996 |
| *Troglodytes hiemalis* | Winter Wren | Troglodytidae | Passeriformes | 460 | 64 | 21.2 | 1893 |
| *Turdus migratorius* | American Robin | Turdidae | Passeriformes | 8649 | 581 | 37.4 | 2996 |
| *Vermivora cyanoptera* | Blue-winged Warbler | Parulidae | Passeriformes | 1015 | 64 | 9.8 | 594 |
| *Vireo gilvus* | Warbling Vireo | Vireonidae | Passeriformes | 2029 | 196 | 25.8 | 2996 |
| *Vireo griseus* | White-eyed Vireo | Vireonidae | Passeriformes | 2649 | 178 | 16.1 | 649 |
| *Vireo olivaceus* | Red-eyed Vireo | Vireonidae | Passeriformes | 2954 | 362 | 27.6 | 1597 |
| *Zonotrichia albicollis* | White-throated Sparrow | Emberizidae | Passeriformes | 2250 | 100 | 16.3 | 1397 |
| *Zonotrichia leucophrys* | White-crowned Sparrow | Emberizidae | Passeriformes | 1147 | 77 | 35.2 | 2996 |

*Table S2*: Change in Size Index and Wing Index per 10 years ($\beta_{IDX}$; Eq 6), 10 degrees latitude ($\gamma_{IDX}$; Eq 11), and 1000 m elevation ($\theta_{IDX}$; Eq 11) for each species.

| Species | $SI\ \beta_{IDX}$ | $SI\ \gamma_{IDX}$ | $SI\ \theta_{IDX}$ | $WI\ \beta_{IDX}$ | $WI\ \gamma_{IDX}$ | $WI\ \theta_{IDX}$ |
|---|---|---|---|---|---|---|
| *Acanthis flammea* | 0.075 | -0.042 | -0.203 | -0.040 | -0.007 | 0.174 |
| *Agelaius phoeniceus* | 0.025 | 0.581 | 0.240 | 0.010 | -0.131 | 0.385 |
| *Ammodramus savannarum* | -0.001 | 0.307 | -0.192 | -0.008 | 0.066 | 0.375 |
| *Baeolophus bicolor* | -0.021 | 0.892 | -0.185 | 0.002 | 0.279 | 0.370 |
| *Bombycilla cedrorum* | -0.071 | 0.216 | -0.038 | 0.109 | -0.088 | 0.052 |
| *Cardellina canadensis* | -0.012 | 0.014 | 0.162 | -0.014 | -0.045 | 0.383 |
| *Cardellina pusilla* | -0.044 | 0.314 | 0.028 | 0.077 | 0.233 | 0.309 |
| *Cardinalis cardinalis* | 0.073 | 0.665 | 0.091 | -0.112 | 0.150 | 0.909 |
| *Catharus fuscescens* | -0.047 | -0.206 | 0.277 | 0.031 | -0.028 | 0.197 |
| *Catharus guttatus* | 0.041 | -0.453 | -0.283 | -0.038 | -0.033 | 0.737 |
| *Catharus ustulatus* | -0.044 | -0.036 | -0.065 | 0.008 | 0.485 | 0.409 |
| *Colaptes auratus* | 0.017 | 0.558 | 0.431 | -0.018 | 0.029 | 0.459 |
| *Dolichonyx oryzivorus* | -0.022 | 0.416 | -0.124 | -0.026 | 0.028 | 0.280 |
| *Dryobates nuttallii* | -0.026 | 0.177 | -0.245 | 0.006 | -0.191 | 0.547 |
| *Dryobates pubescens* | -0.085 | 1.014 | 0.101 | 0.057 | 0.454 | 0.790 |
| *Dryobates villosus* | -0.031 | 1.563 | 0.229 | 0.023 | -0.038 | 0.739 |
| *Dumetella carolinensis* | -0.067 | 0.258 | -0.314 | 0.026 | 0.101 | 0.355 |
| *Empidonax minimus* | -0.048 | 0.237 | -0.097 | 0.117 | 0.061 | 0.207 |
| *Empidonax traillii* | -0.022 | -0.018 | 0.059 | 0.020 | 0.055 | 0.274 |
| *Empidonax virescens* | -0.084 | 0.972 | -0.703 | 0.017 | 0.230 | 0.345 |
| *Geothlypis formosa* | -0.065 | 1.211 | -0.240 | 0.037 | 0.204 | 0.110 |
| *Geothlypis philadelphia* | 0.027 | 0.286 | 0.006 | -0.050 | 0.021 | 0.293 |
| *Geothlypis tolmiei* | -0.061 | 0.809 | 0.273 | -0.002 | 0.088 | 0.324 |
| *Geothlypis trichas* | -0.025 | 0.343 | 0.078 | 0.040 | 0.049 | 0.209 |
| *Haemorhous mexicanus* | 0.020 | 0.542 | -0.348 | -0.034 | -0.095 | 0.447 |
| *Haemorhous purpureus* | -0.066 | 0.946 | -0.426 | 0.063 | 0.135 | 0.366 |
| *Helmitheros vermivorum* | -0.082 | 0.556 | -0.134 | -0.016 | 0.271 | 0.173 |
| *Hylocichla mustelina* | -0.085 | 1.607 | -0.237 | -0.001 | -0.065 | 0.124 |
| *Icteria virens* | -0.065 | 0.530 | -0.100 | -0.044 | 0.162 | 0.594 |
| *Icterus bullockii* | -0.100 | 0.657 | 0.236 | 0.057 | -0.094 | 0.269 |
| *Icterus galbula* | -0.015 | 0.261 | 0.268 | -0.048 | 0.149 | 0.445 |
| *Junco hyemalis* | -0.059 | 0.040 | 0.222 | 0.022 | 0.276 | 0.497 |

*Table S2*: Change in Size Index and Wing Index per 10 years ($\beta_{IDX}$; Eq 6), 10 degrees latitude ($\gamma_{IDX}$; Eq 11), and 1000 m elevation ($\theta_{IDX}$; Eq 11) for each species. *(continued)*

| Species | $SI\ \beta_{IDX}$ | $SI\ \gamma_{IDX}$ | $SI\ \theta_{IDX}$ | $WI\ \beta_{IDX}$ | $WI\ \gamma_{IDX}$ | $WI\ \theta_{IDX}$ |
|---|---|---|---|---|---|---|
| *Limnothlypis swainsonii* | 0.013 | 0.657 | -0.457 | 0.026 | 0.053 | 0.229 |
| *Melospiza georgiana* | -0.024 | -0.247 | -0.260 | -0.016 | 0.155 | 0.334 |
| *Melospiza lincolnii* | -0.055 | -0.463 | 0.137 | 0.005 | -0.248 | 0.091 |
| *Melospiza melodia* | -0.044 | 0.946 | 0.181 | 0.078 | 0.176 | 0.508 |
| *Melozone crissalis* | -0.010 | 1.832 | -0.581 | -0.009 | -0.013 | 0.070 |
| *Mniotilta varia* | -0.056 | -0.037 | 0.235 | 0.069 | -0.235 | 0.207 |
| *Molothrus ater* | 0.033 | 0.723 | -0.239 | 0.055 | 0.140 | 0.342 |
| *Oreothlypis celata* | -0.050 | 0.265 | 0.123 | 0.059 | 0.218 | 0.721 |
| *Oreothlypis luciae* | -0.074 | 0.437 | 0.051 | 0.034 | 0.115 | 0.428 |
| *Oreothlypis peregrina* | -0.033 | -0.142 | -0.037 | 0.087 | 0.056 | 0.398 |
| *Oreothlypis ruficapilla* | -0.048 | 0.131 | 0.172 | 0.032 | 0.064 | 0.152 |
| *Oreothlypis virginiae* | -0.013 | 0.377 | 0.022 | 0.042 | -0.055 | 0.142 |
| *Parkesia motacilla* | -0.028 | -0.171 | -0.262 | -0.008 | 0.282 | 0.310 |
| *Parkesia noveboracensis* | -0.092 | 0.081 | 0.036 | 0.034 | 0.005 | 0.281 |
| *Passerculus sandwichensis* | -0.069 | 0.033 | -0.223 | -0.006 | 0.353 | 0.378 |
| *Passerella iliaca* | 0.012 | 0.265 | -0.371 | -0.010 | 0.277 | 0.391 |
| *Passerina amoena* | 0.019 | 0.008 | -0.099 | 0.021 | 0.102 | 0.007 |
| *Passerina caerulea* | -0.055 | 0.133 | 0.081 | 0.091 | 0.088 | 0.216 |
| *Passerina ciris* | 0.309 | 0.052 | 0.707 | -0.362 | -0.148 | 0.592 |
| *Passerina cyanea* | -0.021 | 0.506 | 0.092 | 0.027 | -0.041 | -0.093 |
| *Pheucticus ludovicianus* | -0.057 | 0.735 | -0.012 | 0.016 | -0.196 | 0.271 |
| *Pheucticus melanocephalus* | -0.022 | 0.425 | 0.268 | 0.047 | 0.224 | 0.254 |
| *Pipilo erythrophthalmus* | -0.043 | -0.288 | -0.115 | 0.104 | 0.550 | 0.253 |
| *Pipilo maculatus* | -0.011 | -0.069 | -0.353 | 0.007 | -0.065 | 0.581 |
| *Piranga ludoviciana* | -0.062 | -0.023 | -0.028 | -0.001 | -0.039 | 0.275 |
| *Piranga olivacea* | -0.042 | 0.514 | -0.097 | 0.045 | 0.116 | 0.053 |
| *Piranga rubra* | 0.034 | -0.294 | 1.091 | -0.067 | 0.135 | 0.456 |
| *Poecile atricapillus* | -0.076 | -0.064 | 0.078 | 0.005 | 0.114 | 0.328 |
| *Poecile carolinensis* | -0.037 | 0.701 | 0.099 | 0.028 | 0.230 | 0.278 |
| *Poecile gambeli* | -0.051 | -0.560 | -0.052 | 0.025 | 0.060 | 0.300 |
| *Poecile rufescens* | -0.018 | 0.191 | -0.196 | 0.067 | 0.227 | 0.471 |
| *Polioptila caerulea* | 0.005 | 0.833 | -0.540 | -0.051 | -0.128 | -0.007 |

*Table S2*: Change in Size Index and Wing Index per 10 years ($\beta_{IDX}$; Eq 6), 10 degrees latitude ($\gamma_{IDX}$; Eq 11), and 1000 m elevation ($\theta_{IDX}$; Eq 11) for each species. *(continued)*

| Species | $SI\ \beta_{IDX}$ | $SI\ \gamma_{IDX}$ | $SI\ \theta_{IDX}$ | $WI\ \beta_{IDX}$ | $WI\ \gamma_{IDX}$ | $WI\ \theta_{IDX}$ |
|---|---|---|---|---|---|---|
| *Protonotaria citrea* | -0.009 | 0.938 | -0.547 | -0.018 | -0.051 | 0.173 |
| *Psaltriparus minimus* | -0.064 | 0.527 | 0.364 | 0.064 | -0.061 | 0.694 |
| *Quiscalus quiscula* | -0.009 | 0.435 | -0.388 | 0.029 | 0.050 | 0.290 |
| *Regulus calendula* | 0.011 | 0.230 | 0.046 | 0.052 | -0.204 | 0.553 |
| *Regulus satrapa* | -0.089 | 0.000 | -0.252 | 0.057 | -0.161 | 0.101 |
| *Seiurus aurocapilla* | -0.051 | 0.280 | -0.293 | 0.045 | -0.002 | 0.180 |
| *Setophaga americana* | -0.072 | 0.472 | -0.126 | 0.091 | 0.252 | 0.345 |
| *Setophaga citrina* | -0.002 | 0.994 | -0.232 | 0.029 | 0.266 | 0.383 |
| *Setophaga coronata* | -0.115 | 0.338 | 0.449 | -0.057 | -0.067 | 0.559 |
| *Setophaga discolor* | -0.021 | 0.704 | -0.645 | 0.007 | -0.028 | 0.256 |
| *Setophaga magnolia* | -0.012 | -0.106 | -0.121 | 0.043 | -0.078 | 0.357 |
| *Setophaga occidentalis* | -0.069 | 1.396 | -0.115 | 0.077 | -0.048 | 0.106 |
| *Setophaga pensylvanica* | -0.077 | -0.075 | -0.054 | 0.035 | -0.099 | 0.132 |
| *Setophaga petechia* | -0.055 | 0.166 | -0.327 | 0.098 | -0.073 | 0.095 |
| *Setophaga ruticilla* | -0.073 | 0.014 | -0.007 | 0.112 | -0.307 | -0.047 |
| *Setophaga townsendi* | -0.078 | 0.278 | -0.177 | 0.045 | 0.058 | 0.279 |
| *Setophaga virens* | 0.005 | 0.574 | 0.098 | 0.028 | 0.013 | 0.411 |
| *Sitta canadensis* | -0.052 | 0.095 | -0.134 | 0.102 | -0.069 | 0.161 |
| *Sitta carolinensis* | -0.018 | 0.588 | -0.423 | 0.028 | -0.179 | 0.567 |
| *Sphyrapicus nuchalis* | 0.048 | 0.217 | -0.193 | -0.019 | -0.096 | 0.215 |
| *Sphyrapicus varius* | -0.061 | 0.339 | -0.469 | 0.004 | -0.083 | 0.239 |
| *Spinus pinus* | 0.010 | 0.282 | -0.149 | -0.037 | -0.046 | 0.401 |
| *Spinus psaltria* | 0.012 | 0.059 | -0.229 | -0.010 | -0.122 | 0.287 |
| *Spinus tristis* | 0.000 | 0.755 | 0.257 | 0.012 | -0.095 | 0.287 |
| *Spiza americana* | 0.008 | 0.431 | 0.127 | 0.035 | -0.147 | 0.180 |
| *Spizella pallida* | -0.098 | 0.225 | 0.185 | 0.043 | -0.033 | 0.223 |
| *Spizella passerina* | -0.057 | 0.281 | 0.274 | 0.083 | 0.267 | 0.423 |
| *Spizella pusilla* | -0.045 | 0.219 | 0.223 | -0.094 | -0.461 | 0.253 |
| *Tachycineta bicolor* | -0.118 | 0.804 | -0.163 | 0.037 | 0.041 | 0.367 |
| *Thryomanes bewickii* | -0.042 | 0.277 | -0.045 | -0.023 | -0.523 | 0.552 |
| *Thryothorus ludovicianus* | 0.036 | 0.558 | -0.402 | -0.012 | -0.041 | 0.453 |

*Table S2*: Change in Size Index and Wing Index per 10 years ($\beta_{IDX}$; Eq 6), 10 degrees latitude ($\gamma_{IDX}$; Eq 11), and 1000 m elevation ($\theta_{IDX}$; Eq 11) for each species. *(continued)*

| Species | $SI\ \beta_{IDX}$ | $SI\ \gamma_{IDX}$ | $SI\ \theta_{IDX}$ | $WI\ \beta_{IDX}$ | $WI\ \gamma_{IDX}$ | $WI\ \theta_{IDX}$ |
|---|---|---|---|---|---|---|
| *Toxostoma rufum* | -0.024 | 0.442 | 0.095 | -0.050 | -0.014 | 0.230 |
| *Troglodytes aedon* | 0.072 | 0.396 | -0.104 | -0.001 | -0.118 | 0.277 |
| *Troglodytes hiemalis* | -0.056 | 0.276 | -0.474 | 0.104 | 0.095 | 0.205 |
| *Turdus migratorius* | -0.058 | 0.364 | 0.272 | -0.001 | 0.030 | 0.490 |
| *Vermivora cyanoptera* | 0.019 | 1.247 | -0.315 | 0.028 | -0.118 | 0.209 |
| *Vireo gilvus* | -0.051 | 0.194 | -0.189 | 0.016 | -0.208 | 0.372 |
| *Vireo griseus* | 0.068 | 0.614 | -0.575 | -0.021 | 0.254 | 0.419 |
| *Vireo olivaceus* | -0.022 | 0.439 | -0.208 | -0.048 | 0.001 | 0.169 |
| *Zonotrichia albicollis* | -0.056 | 0.281 | -0.188 | 0.026 | -0.025 | 0.153 |
| *Zonotrichia leucophrys* | 0.038 | -0.241 | 0.288 | 0.067 | 0.621 | 0.632 |

*Table S3*: Percent change in mass and wing length over 30 year period of study ($\omega_{M_{TIME}}$ and $\omega_{W_{TIME}}$, respectively; Eq 25) and across the latitudal ($\omega_{M_{LAT}}$ and $\omega_{W_{LAT}}$, respectively; Eq 25) and elevational range ($\omega_{M_{ELEV}}$ and $\omega_{W_{ELEV}}$, respectively; Eq 25) sampled for each species.

| Species | $\omega_{M_{TIME}}$ | $\omega_{M_{LAT}}$ | $\omega_{M_{ELEV}}$ | $\omega_{W_{TIME}}$ | $\omega_{W_{LAT}}$ | $\omega_{W_{ELEV}}$ |
|---|---|---|---|---|---|---|
| *Acanthis flammea* | 1.521 | -0.355 | -1.440 | 0.152 | -0.149 | 0.045 |
| *Agelaius phoeniceus* | 0.560 | 19.908 | 3.777 | 0.307 | 4.230 | 5.085 |
| *Ammodramus savannarum* | 0.010 | 0.877 | -0.554 | -0.077 | 0.402 | 0.287 |
| *Baeolophus bicolor* | -0.510 | 12.262 | -2.174 | -0.148 | 5.908 | 0.968 |
| *Bombycilla cedrorum* | -1.797 | 3.629 | -0.681 | 0.471 | 0.542 | 0.143 |
| *Cardellina canadensis* | -0.180 | 0.283 | 1.003 | -0.191 | -0.215 | 2.288 |
| *Cardellina pusilla* | -1.237 | 7.321 | -0.426 | 0.465 | 5.477 | 3.418 |
| *Cardinalis cardinalis* | 1.942 | 9.444 | -0.440 | -0.540 | 4.128 | 4.535 |
| *Catharus fuscescens* | -0.962 | -2.121 | 3.735 | -0.002 | -0.877 | 2.936 |
| *Catharus guttatus* | 1.775 | -15.411 | -12.408 | -0.007 | -5.916 | 6.513 |
| *Catharus ustulatus* | -0.758 | -2.512 | -2.378 | -0.172 | 5.410 | 3.635 |
| *Colaptes auratus* | 0.646 | 18.681 | 11.765 | -0.030 | 6.262 | 9.749 |
| *Dolichonyx oryzivorus* | -0.279 | 0.764 | -0.561 | -0.368 | 0.286 | 0.381 |
| *Dryobates nuttallii* | -0.513 | 1.022 | -1.897 | -0.120 | -0.076 | 0.805 |
| *Dryobates pubescens* | -2.151 | 27.492 | -0.196 | -0.097 | 14.468 | 7.734 |
| *Dryobates villosus* | -0.962 | 58.304 | 3.510 | -0.028 | 15.961 | 10.509 |
| *Dumetella carolinensis* | -1.182 | 3.056 | -4.988 | -0.127 | 1.824 | 1.322 |
| *Empidonax minimus* | -1.178 | 2.660 | -1.166 | 0.688 | 1.234 | 0.579 |
| *Empidonax traillii* | -0.566 | -0.343 | 0.385 | 0.047 | 0.278 | 2.882 |
| *Empidonax virescens* | -1.931 | 9.599 | -3.433 | -0.442 | 4.369 | -0.304 |
| *Geothlypis formosa* | -1.509 | 9.258 | -0.939 | -0.054 | 3.918 | -0.081 |
| *Geothlypis philadelphia* | 0.663 | 2.815 | -0.157 | -0.275 | 1.036 | 0.575 |
| *Geothlypis tolmiei* | -1.150 | 10.415 | 4.292 | -0.405 | 3.983 | 4.944 |
| *Geothlypis trichas* | -0.679 | 8.142 | 0.799 | 0.258 | 3.306 | 2.425 |
| *Haemorhous mexicanus* | 0.504 | 8.074 | -5.984 | -0.170 | 1.959 | 1.216 |
| *Haemorhous purpureus* | -1.555 | 16.255 | -7.482 | 0.182 | 6.396 | 0.567 |
| *Helmitheros vermivorum* | -1.559 | 3.281 | -0.802 | -0.695 | 2.006 | 0.195 |
| *Hylocichla mustelina* | -1.680 | 18.882 | -1.913 | -0.569 | 5.562 | -0.165 |
| *Icteria virens* | -1.200 | 8.903 | -3.085 | -0.955 | 4.587 | 4.454 |
| *Icterus bullockii* | -2.686 | 12.625 | 3.504 | -0.191 | 3.179 | 3.573 |
| *Icterus galbula* | -0.094 | 2.964 | 1.442 | -0.598 | 2.228 | 2.815 |
| *Junco hyemalis* | -1.243 | -0.167 | 2.728 | -0.165 | 3.606 | 6.899 |

*Table S3*: Percent change in mass and wing length over 30 year period of study ($\omega_{M_{TIME}}$ and $\omega_{W_{TIME}}$, respectively; Eq 25) and across the latitudal ($\omega_{M_{LAT}}$ and $\omega_{W_{LAT}}$, respectively; Eq 25) and elevational range ($\omega_{M_{ELEV}}$ and $\omega_{W_{ELEV}}$, respectively; Eq 25) sampled for each species. *(continued)*

| Species | $\omega_{M_{TIME}}$ | $\omega_{M_{LAT}}$ | $\omega_{M_{ELEV}}$ | $\omega_{W_{TIME}}$ | $\omega_{W_{LAT}}$ | $\omega_{W_{ELEV}}$ |
|---|---|---|---|---|---|---|
| *Limnothlypis swainsonii* | 0.160 | 2.486 | -0.296 | 0.294 | 0.927 | -0.027 |
| *Melospiza georgiana* | -0.491 | -3.389 | -2.792 | -0.360 | -0.106 | 0.669 |
| *Melospiza lincolnii* | -1.174 | -9.196 | 2.640 | -0.337 | -6.035 | 1.906 |
| *Melospiza melodia* | -1.414 | 22.552 | 1.986 | 0.612 | 9.397 | 6.825 |
| *Melozone crissalis* | -0.277 | 21.775 | -9.787 | -0.202 | 6.724 | -2.939 |
| *Mniotilta varia* | -1.302 | 0.069 | 2.111 | 0.325 | -2.300 | 1.928 |
| *Molothrus ater* | 0.773 | 20.034 | -6.471 | 0.856 | 7.733 | 0.892 |
| *Oreothlypis celata* | -1.274 | 5.670 | -0.266 | 0.361 | 5.379 | 9.098 |
| *Oreothlypis luciae* | -1.654 | 1.733 | -0.260 | -0.124 | 0.861 | 2.663 |
| *Oreothlypis peregrina* | -0.835 | -1.101 | -0.692 | 0.556 | -0.148 | 1.256 |
| *Oreothlypis ruficapilla* | -1.019 | 1.323 | 1.802 | -0.003 | 0.821 | 1.615 |
| *Oreothlypis virginiae* | -0.382 | 1.651 | -0.013 | 0.356 | 0.408 | 0.431 |
| *Parkesia motacilla* | -0.555 | -1.795 | -1.383 | -0.268 | 0.557 | 0.209 |
| *Parkesia noveboracensis* | -1.927 | 1.532 | -0.106 | -0.289 | 0.560 | 2.384 |
| *Passerculus sandwichensis* | -1.476 | -0.416 | -5.376 | -0.560 | 3.953 | 1.906 |
| *Passerella iliaca* | 0.503 | 11.163 | -14.143 | 0.043 | 7.859 | -0.228 |
| *Passerina amoena* | 0.331 | -0.063 | -1.408 | 0.309 | 0.511 | -0.427 |
| *Passerina caerulea* | -1.333 | 1.241 | 0.552 | 0.503 | 0.897 | 1.586 |
| *Passerina ciris* | 11.037 | 0.626 | 3.577 | -1.219 | -0.358 | 2.681 |
| *Passerina cyanea* | -0.481 | 5.618 | 1.103 | 0.111 | 1.611 | -0.126 |
| *Pheucticus ludovicianus* | -1.174 | 11.387 | -0.581 | -0.210 | 2.061 | 1.398 |
| *Pheucticus melanocephalus* | -0.674 | 6.335 | 5.199 | 0.308 | 3.791 | 4.441 |
| *Pipilo erythrophthalmus* | -1.298 | -4.377 | -1.777 | 0.884 | 2.343 | 1.108 |
| *Pipilo maculatus* | -0.237 | -0.749 | -6.814 | 0.015 | -0.829 | 3.551 |
| *Piranga ludoviciana* | -1.214 | -0.271 | -1.319 | -0.414 | -0.446 | 2.292 |
| *Piranga olivacea* | -0.970 | 5.074 | -0.705 | 0.158 | 2.309 | -0.038 |
| *Piranga rubra* | 1.174 | -3.165 | 15.288 | -0.388 | -0.476 | 7.686 |
| *Poecile atricapillus* | -1.641 | -1.620 | 0.434 | -0.492 | 0.723 | 3.531 |
| *Poecile carolinensis* | -1.216 | 8.684 | 0.689 | -0.037 | 4.111 | 1.525 |
| *Poecile gambeli* | -1.023 | -5.849 | -1.367 | -0.108 | -1.674 | 1.716 |
| *Poecile rufescens* | -0.504 | 1.859 | -2.405 | 0.545 | 2.504 | 1.815 |
| *Polioptila caerulea* | 0.348 | 10.870 | -11.077 | -0.512 | 2.776 | -3.908 |

*Table S3*: Percent change in mass and wing length over 30 year period of study ($\omega_{M_{TIME}}$ and $\omega_{W_{TIME}}$, respectively; Eq 25) and across the latitudal ($\omega_{M_{LAT}}$ and $\omega_{W_{LAT}}$, respectively; Eq 25) and elevational range ($\omega_{M_{ELEV}}$ and $\omega_{W_{ELEV}}$, respectively; Eq 25) sampled for each species. *(continued)*

| Species | $\omega_{M_{TIME}}$ | $\omega_{M_{LAT}}$ | $\omega_{M_{ELEV}}$ | $\omega_{W_{TIME}}$ | $\omega_{W_{LAT}}$ | $\omega_{W_{ELEV}}$ |
|---|---|---|---|---|---|---|
| *Protonotaria citrea* | -0.122 | 9.645 | -1.074 | -0.232 | 2.852 | -0.193 |
| *Psaltriparus minimus* | -1.665 | 7.129 | 4.438 | 0.257 | 1.876 | 8.689 |
| *Quiscalus quiscula* | -0.322 | 9.832 | -6.657 | 0.217 | 3.647 | -0.278 |
| *Regulus calendula* | -0.006 | 6.389 | -1.565 | 0.795 | -1.379 | 8.157 |
| *Regulus satrapa* | -2.108 | 0.539 | -4.636 | -0.039 | -1.501 | -0.615 |
| *Seiurus aurocapilla* | -1.078 | 4.204 | -3.046 | 0.111 | 1.365 | -0.057 |
| *Setophaga americana* | -1.900 | 6.365 | -0.862 | 0.425 | 4.018 | 0.591 |
| *Setophaga citrina* | -0.139 | 9.167 | -2.125 | 0.271 | 4.265 | 0.734 |
| *Setophaga coronata* | -1.980 | 7.860 | 6.831 | -1.338 | 1.637 | 9.285 |
| *Setophaga discolor* | -0.471 | 5.445 | -2.939 | -0.084 | 1.674 | -0.428 |
| *Setophaga magnolia* | -0.339 | -0.873 | -1.317 | 0.349 | -0.740 | 1.109 |
| *Setophaga occidentalis* | -1.649 | 7.858 | -1.989 | 0.222 | 2.424 | 0.122 |
| *Setophaga pensylvanica* | -1.729 | -0.697 | -0.823 | -0.185 | -0.891 | 0.524 |
| *Setophaga petechia* | -1.627 | 5.472 | -7.792 | 0.590 | 0.709 | -1.593 |
| *Setophaga ruticilla* | -1.938 | 1.272 | 0.017 | 0.663 | -2.793 | -0.250 |
| *Setophaga townsendi* | -1.669 | 2.827 | -2.235 | -0.099 | 1.245 | 0.744 |
| *Setophaga virens* | 0.012 | 6.922 | 0.148 | 0.274 | 2.344 | 1.308 |
| *Sitta canadensis* | -1.204 | 1.714 | -2.566 | 0.622 | -0.059 | 0.638 |
| *Sitta carolinensis* | -0.685 | 14.161 | -13.943 | 0.124 | 2.949 | 1.757 |
| *Sphyrapicus nuchalis* | 1.211 | 3.099 | -4.235 | 0.209 | 0.461 | 0.417 |
| *Sphyrapicus varius* | -1.251 | 5.245 | -3.907 | -0.384 | 1.189 | -0.501 |
| *Spinus pinus* | 0.347 | 6.593 | -4.505 | -0.287 | 1.642 | 2.880 |
| *Spinus psaltria* | 0.277 | 0.554 | -3.803 | -0.005 | -0.244 | 0.739 |
| *Spinus tristis* | -0.032 | 13.173 | 3.531 | 0.116 | 3.374 | 3.734 |
| *Spiza americana* | 0.044 | 3.237 | 0.363 | 0.406 | 0.486 | 0.496 |
| *Spizella pallida* | -1.877 | 1.682 | 0.993 | -0.201 | 0.424 | 1.160 |
| *Spizella passerina* | -1.295 | 4.977 | 3.339 | 0.486 | 5.185 | 5.603 |
| *Spizella pusilla* | -0.580 | 3.395 | 0.795 | -1.277 | -1.811 | 0.898 |
| *Tachycineta bicolor* | -2.778 | 22.852 | -3.336 | -0.547 | 7.613 | 1.560 |
| *Thryomanes bewickii* | -0.968 | 7.412 | -2.598 | -0.687 | -3.690 | 5.395 |
| *Thryothorus ludovicianus* | 0.922 | 8.206 | -2.402 | 0.167 | 2.397 | 0.280 |

*Table S3*: Percent change in mass and wing length over 30 year period of study ($\omega_{M_{TIME}}$ and $\omega_{W_{TIME}}$, respectively; Eq 25) and across the latitudal ($\omega_{M_{LAT}}$ and $\omega_{W_{LAT}}$, respectively; Eq 25) and elevational range ($\omega_{M_{ELEV}}$ and $\omega_{W_{ELEV}}$, respectively; Eq 25) sampled for each species. *(continued)*

| Species | $\omega_{M_{TIME}}$ | $\omega_{M_{LAT}}$ | $\omega_{M_{ELEV}}$ | $\omega_{W_{TIME}}$ | $\omega_{W_{LAT}}$ | $\omega_{W_{ELEV}}$ |
|---|---|---|---|---|---|---|
| *Toxostoma rufum* | -0.262 | 6.434 | 0.318 | -0.695 | 1.978 | 0.996 |
| *Troglodytes aedon* | 1.598 | 7.566 | -3.290 | 0.521 | 1.216 | 2.519 |
| *Troglodytes hiemalis* | -1.481 | 3.611 | -6.091 | 0.881 | 2.089 | -0.389 |
| *Turdus migratorius* | -1.240 | 10.102 | 4.220 | -0.424 | 3.706 | 7.302 |
| *Vermivora cyanoptera* | 0.324 | 9.720 | -1.517 | 0.431 | 2.687 | -0.042 |
| *Vireo gilvus* | -1.081 | 4.139 | -5.013 | -0.172 | -0.759 | 2.714 |
| *Vireo griseus* | 1.603 | 7.067 | -3.053 | 0.266 | 4.059 | 0.096 |
| *Vireo olivaceus* | -0.259 | 7.485 | -2.205 | -0.540 | 2.444 | 0.108 |
| *Zonotrichia albicollis* | -1.002 | 2.634 | -1.641 | -0.111 | 0.752 | 0.057 |
| *Zonotrichia leucophrys* | 0.695 | -9.935 | 4.823 | 1.211 | 7.319 | 11.317 |

*Table S4*: Change in SI per 1°C change in June max temperature ($\gamma_{TVT}$; Eq 15) and the change in this effect per 10°C change in mean June max temperature over space ($\theta_{TVT}$; Eq 15) for each species. The effect of temperature in the year of capture (lag-0), as well as temperature one (lag-1) and two years (lag-2) prior to capture is presented.

| Species | $\gamma_{TVT}$ - lag-0 | $\gamma_{TVT}$ - lag-1 | $\gamma_{TVT}$ - lag-2 | $\theta_{TVT}$ - lag-0 | $\theta_{TVT}$ - lag-1 | $\theta_{TVT}$ - lag-2 |
|---|---|---|---|---|---|---|
| *Acanthis flammea* | -0.010 | -0.004 | 0.001 | -0.011 | 0.019 | 0.037 |
| *Agelaius phoeniceus* | -0.012 | -0.007 | 0.001 | 0.051 | 0.046 | 0.044 |
| *Ammodramus savannarum* | -0.006 | 0.007 | -0.006 | -0.031 | -0.016 | -0.032 |
| *Baeolophus bicolor* | -0.006 | -0.004 | -0.002 | -0.041 | -0.047 | -0.045 |
| *Bombycilla cedrorum* | -0.011 | 0.005 | -0.002 | 0.009 | 0.011 | 0.013 |
| *Cardellina canadensis* | -0.009 | -0.002 | 0.001 | -0.019 | -0.009 | -0.023 |
| *Cardellina pusilla* | -0.009 | -0.019 | -0.001 | 0.012 | 0.017 | 0.016 |
| *Cardinalis cardinalis* | -0.008 | -0.006 | 0.009 | 0.019 | -0.021 | 0.033 |
| *Catharus fuscescens* | -0.010 | -0.004 | -0.001 | -0.010 | -0.034 | -0.017 |
| *Catharus guttatus* | -0.011 | 0.004 | -0.008 | -0.014 | -0.028 | -0.028 |
| *Catharus ustulatus* | -0.010 | -0.010 | 0.004 | -0.047 | -0.010 | -0.008 |
| *Colaptes auratus* | -0.014 | -0.009 | 0.002 | 0.046 | 0.040 | 0.058 |
| *Dolichonyx oryzivorus* | -0.010 | -0.002 | -0.001 | -0.012 | -0.006 | -0.008 |
| *Dryobates nuttallii* | -0.017 | -0.010 | 0.001 | 0.025 | 0.025 | 0.025 |
| *Dryobates pubescens* | -0.015 | -0.014 | -0.004 | -0.052 | -0.050 | -0.056 |
| *Dryobates villosus* | -0.015 | -0.007 | -0.001 | -0.002 | -0.030 | -0.013 |
| *Dumetella carolinensis* | -0.015 | -0.004 | -0.005 | -0.056 | -0.024 | -0.076 |
| *Empidonax minimus* | -0.006 | 0.003 | 0.002 | -0.039 | -0.037 | 0.006 |
| *Empidonax traillii* | -0.010 | -0.005 | -0.002 | -0.035 | -0.031 | -0.029 |
| *Empidonax virescens* | -0.009 | -0.004 | -0.001 | -0.053 | -0.038 | -0.038 |
| *Geothlypis formosa* | -0.012 | -0.015 | -0.004 | -0.064 | -0.041 | 0.026 |
| *Geothlypis philadelphia* | -0.010 | 0.001 | 0.000 | -0.020 | -0.014 | 0.006 |
| *Geothlypis tolmiei* | -0.018 | -0.006 | -0.003 | 0.011 | 0.019 | -0.012 |
| *Geothlypis trichas* | -0.015 | 0.001 | 0.000 | 0.015 | 0.009 | 0.011 |
| *Haemorhous mexicanus* | -0.003 | -0.009 | -0.001 | -0.014 | -0.021 | -0.006 |
| *Haemorhous purpureus* | -0.017 | -0.018 | -0.005 | 0.000 | 0.013 | 0.023 |
| *Helmitheros vermivorum* | -0.014 | -0.007 | -0.005 | -0.019 | -0.022 | -0.030 |
| *Hylocichla mustelina* | -0.014 | -0.006 | -0.008 | -0.083 | -0.077 | -0.105 |
| *Icteria virens* | -0.009 | -0.003 | -0.001 | -0.005 | -0.024 | -0.011 |
| *Icterus bullockii* | -0.012 | -0.007 | -0.001 | -0.015 | -0.010 | -0.016 |
| *Icterus galbula* | -0.015 | -0.003 | -0.001 | 0.012 | 0.017 | 0.012 |

*Table S4*: Change in SI per 1°C change in June max temperature ($\gamma_{TVT}$; Eq 15) and the change in this effect per 10°C change in mean June max temperature over space ($\theta_{TVT}$; Eq 15) for each species. The effect of temperature in the year of capture (lag-0), as well as temperature one (lag-1) and two years (lag-2) prior to capture is presented. *(continued)*

| Species | $\gamma_{TVT}$ - lag-0 | $\gamma_{TVT}$ - lag-1 | $\gamma_{TVT}$ - lag-2 | $\theta_{TVT}$ - lag-0 | $\theta_{TVT}$ - lag-1 | $\theta_{TVT}$ - lag-2 |
|---|---|---|---|---|---|---|
| *Junco hyemalis* | -0.013 | -0.002 | -0.005 | 0.003 | 0.015 | 0.001 |
| *Limnothlypis swainsonii* | -0.011 | -0.001 | -0.001 | -0.020 | -0.015 | -0.012 |
| *Melospiza georgiana* | -0.008 | -0.002 | -0.005 | -0.029 | -0.033 | -0.054 |
| *Melospiza lincolnii* | -0.002 | -0.003 | 0.003 | -0.005 | 0.032 | 0.014 |
| *Melospiza melodia* | -0.011 | 0.007 | -0.003 | -0.003 | -0.009 | -0.006 |
| *Melozone crissalis* | -0.016 | 0.001 | 0.005 | 0.090 | 0.101 | 0.100 |
| *Mniotilta varia* | -0.010 | 0.003 | -0.002 | -0.017 | -0.017 | -0.006 |
| *Molothrus ater* | -0.001 | -0.004 | -0.005 | -0.093 | -0.087 | -0.092 |
| *Oreothlypis celata* | -0.009 | -0.007 | 0.004 | 0.036 | 0.025 | 0.045 |
| *Oreothlypis luciae* | -0.014 | -0.009 | -0.001 | 0.009 | -0.001 | -0.003 |
| *Oreothlypis peregrina* | -0.015 | 0.001 | -0.002 | 0.004 | -0.001 | -0.007 |
| *Oreothlypis ruficapilla* | -0.005 | 0.001 | -0.001 | -0.007 | 0.024 | -0.009 |
| *Oreothlypis virginiae* | -0.012 | -0.004 | 0.000 | 0.003 | -0.015 | 0.008 |
| *Parkesia motacilla* | -0.010 | -0.005 | -0.001 | -0.022 | -0.014 | 0.009 |
| *Parkesia noveboracensis* | -0.009 | -0.002 | -0.001 | -0.022 | -0.030 | -0.035 |
| *Passerculus sandwichensis* | -0.012 | -0.003 | 0.002 | 0.001 | 0.011 | 0.053 |
| *Passerella iliaca* | -0.015 | -0.005 | 0.001 | 0.037 | 0.013 | 0.036 |
| *Passerina amoena* | -0.010 | 0.004 | -0.004 | 0.004 | 0.001 | 0.007 |
| *Passerina caerulea* | -0.011 | -0.007 | -0.001 | -0.020 | -0.025 | -0.006 |
| *Passerina ciris* | 0.000 | 0.000 | 0.002 | -0.015 | 0.012 | 0.005 |
| *Passerina cyanea* | -0.014 | 0.009 | 0.001 | 0.004 | -0.002 | 0.004 |
| *Pheucticus ludovicianus* | -0.015 | -0.004 | 0.000 | -0.024 | -0.001 | 0.007 |
| *Pheucticus melanocephalus* | -0.020 | -0.002 | 0.002 | 0.009 | 0.009 | 0.011 |
| *Pipilo erythrophthalmus* | -0.010 | -0.001 | 0.002 | 0.018 | 0.021 | 0.025 |
| *Pipilo maculatus* | -0.021 | -0.015 | 0.001 | 0.049 | 0.038 | 0.038 |
| *Piranga ludoviciana* | -0.013 | -0.003 | 0.001 | -0.004 | -0.013 | 0.001 |
| *Piranga olivacea* | -0.011 | -0.008 | 0.000 | -0.019 | -0.007 | -0.002 |
| *Piranga rubra* | -0.018 | -0.015 | -0.006 | 0.026 | 0.036 | 0.014 |
| *Poecile atricapillus* | -0.009 | 0.002 | 0.000 | -0.010 | -0.008 | -0.001 |
| *Poecile carolinensis* | -0.011 | -0.013 | -0.006 | -0.047 | -0.045 | -0.037 |
| *Poecile gambeli* | -0.010 | -0.005 | -0.004 | -0.050 | -0.023 | -0.043 |

*Table S4*: Change in SI per 1°C change in June max temperature ($\gamma_{TVT}$; Eq 15) and the change in this effect per 10°C change in mean June max temperature over space ($\theta_{TVT}$; Eq 15) for each species. The effect of temperature in the year of capture (lag-0), as well as temperature one (lag-1) and two years (lag-2) prior to capture is presented. *(continued)*

| Species | $\gamma_{TVT}$ - lag-0 | $\gamma_{TVT}$ - lag-1 | $\gamma_{TVT}$ - lag-2 | $\theta_{TVT}$ - lag-0 | $\theta_{TVT}$ - lag-1 | $\theta_{TVT}$ - lag-2 |
|---|---|---|---|---|---|---|
| *Poecile rufescens* | -0.010 | -0.006 | -0.001 | -0.019 | -0.006 | -0.003 |
| *Polioptila caerulea* | -0.006 | 0.002 | -0.002 | -0.049 | -0.078 | -0.045 |
| *Protonotaria citrea* | -0.013 | -0.002 | 0.004 | -0.025 | -0.031 | 0.036 |
| *Psaltriparus minimus* | -0.011 | -0.006 | 0.002 | 0.001 | 0.011 | 0.018 |
| *Quiscalus quiscula* | -0.010 | 0.003 | -0.002 | -0.049 | -0.016 | -0.029 |
| *Regulus calendula* | -0.010 | -0.006 | 0.000 | -0.001 | -0.004 | 0.014 |
| *Regulus satrapa* | -0.015 | -0.005 | -0.004 | -0.006 | -0.012 | -0.022 |
| *Seiurus aurocapilla* | -0.017 | -0.004 | -0.002 | -0.044 | -0.033 | -0.042 |
| *Setophaga americana* | -0.014 | -0.009 | -0.001 | -0.018 | -0.013 | -0.001 |
| *Setophaga citrina* | -0.007 | 0.004 | 0.006 | -0.043 | -0.022 | 0.033 |
| *Setophaga coronata* | -0.010 | 0.003 | -0.001 | 0.034 | 0.047 | 0.031 |
| *Setophaga discolor* | -0.011 | -0.003 | -0.001 | -0.002 | 0.035 | 0.009 |
| *Setophaga magnolia* | -0.014 | -0.003 | 0.004 | 0.024 | 0.025 | 0.031 |
| *Setophaga occidentalis* | -0.018 | -0.016 | -0.003 | 0.018 | 0.020 | 0.019 |
| *Setophaga pensylvanica* | -0.009 | 0.001 | 0.000 | -0.033 | -0.030 | 0.009 |
| *Setophaga petechia* | -0.016 | -0.013 | 0.004 | -0.012 | -0.013 | -0.008 |
| *Setophaga ruticilla* | -0.014 | -0.007 | -0.002 | -0.019 | -0.013 | -0.025 |
| *Setophaga townsendi* | -0.010 | -0.008 | -0.003 | 0.005 | 0.004 | -0.005 |
| *Setophaga virens* | -0.012 | -0.004 | -0.002 | -0.014 | -0.007 | -0.002 |
| *Sitta canadensis* | -0.010 | -0.004 | -0.001 | -0.004 | 0.008 | 0.007 |
| *Sitta carolinensis* | -0.004 | -0.002 | -0.009 | -0.119 | -0.106 | -0.129 |
| *Sphyrapicus nuchalis* | -0.007 | 0.001 | -0.001 | -0.026 | 0.007 | -0.011 |
| *Sphyrapicus varius* | -0.014 | -0.006 | -0.002 | 0.000 | 0.004 | -0.009 |
| *Spinus pinus* | -0.016 | 0.000 | 0.004 | 0.016 | -0.010 | 0.032 |
| *Spinus psaltria* | -0.014 | 0.000 | 0.000 | 0.012 | -0.001 | 0.011 |
| *Spinus tristis* | -0.013 | -0.004 | -0.007 | -0.063 | -0.055 | -0.066 |
| *Spiza americana* | -0.017 | -0.001 | -0.001 | 0.018 | 0.006 | -0.008 |
| *Spizella pallida* | -0.016 | -0.008 | 0.001 | 0.001 | 0.001 | 0.018 |
| *Spizella passerina* | -0.016 | -0.005 | -0.002 | -0.020 | -0.016 | -0.042 |
| *Spizella pusilla* | -0.010 | -0.002 | -0.002 | 0.002 | -0.002 | -0.007 |
| *Tachycineta bicolor* | -0.014 | -0.009 | 0.000 | -0.004 | -0.009 | -0.005 |

*Table S4*: Change in SI per 1°C change in June max temperature ($\gamma_{TVT}$; Eq 15) and the change in this effect per 10°C change in mean June max temperature over space ($\theta_{TVT}$; Eq 15) for each species. The effect of temperature in the year of capture (lag-0), as well as temperature one (lag-1) and two years (lag-2) prior to capture is presented. *(continued)*

| Species | $\gamma_{TVT}$ - lag-0 | $\gamma_{TVT}$ - lag-1 | $\gamma_{TVT}$ - lag-2 | $\theta_{TVT}$ - lag-0 | $\theta_{TVT}$ - lag-1 | $\theta_{TVT}$ - lag-2 |
|---|---|---|---|---|---|---|
| *Thryomanes bewickii* | -0.013 | -0.007 | 0.005 | 0.047 | 0.041 | 0.043 |
| *Thryothorus ludovicianus* | -0.008 | 0.001 | -0.001 | -0.028 | -0.049 | -0.010 |
| *Toxostoma rufum* | -0.013 | -0.012 | 0.000 | 0.022 | 0.030 | 0.019 |
| *Troglodytes aedon* | -0.012 | 0.002 | 0.004 | -0.045 | -0.043 | -0.029 |
| *Troglodytes hiemalis* | -0.008 | -0.002 | 0.000 | -0.020 | -0.020 | -0.014 |
| *Turdus migratorius* | -0.025 | -0.003 | -0.001 | 0.000 | 0.005 | 0.002 |
| *Vermivora cyanoptera* | -0.017 | -0.005 | 0.000 | -0.003 | -0.004 | -0.007 |
| *Vireo gilvus* | -0.022 | -0.008 | 0.000 | 0.028 | 0.026 | 0.016 |
| *Vireo griseus* | -0.003 | -0.001 | -0.001 | -0.058 | -0.014 | -0.025 |
| *Vireo olivaceus* | -0.011 | -0.010 | -0.004 | -0.033 | 0.000 | -0.027 |
| *Zonotrichia albicollis* | -0.011 | 0.000 | -0.001 | -0.036 | -0.015 | -0.030 |
| *Zonotrichia leucophrys* | -0.008 | -0.003 | 0.004 | -0.030 | -0.035 | -0.006 |
